# Subcellular Cavitation Bubbles Induce Cellular Mechanolysis and Collective Wound Healing in Ultrasound-Inflicted Cell Ablation

**DOI:** 10.1101/2023.11.20.567805

**Authors:** Ziyue Bai, Zaimeng Li, Yue Shao

## Abstract

Focused ultrasound (FUS) has been widely adopted in medical and life science researches. Although various physical and biological effects of FUS have been well-documented, there is still a lack of understanding and direct evidence on the biological mechanism of therapeutic cell ablation caused by high-intensity ultrasound (HIFU) and the subsequent wound healing responses. This study develops an enclosed cell culture device that synergistically combines non-invasive FUS stimulation and real-time, on-the-fly live-cell imaging, providing an in vitro platform to explore short and long-term biological effects of ultrasound. The process, mechanism, and wound healing response of cell ablation induced by HIFU are elucidated, revealing a unique mechanism, termed ultrasound-inflicted cellular mechanolysis, that is mediated by growing subcellular cavitation air bubbles under confined contact with cells. This provides a previously unappreciated mechanism for understanding the biomechanical principles of ultrasound-based ablative therapy. We also reveal a post-ablation phantom layer that serves as a guiding cue for collective cell migration during wound healing, thereby providing a biomimetic model for studying wound healing after HIFU-inflicted damage. Together, this study provides theoretical and technological basis for advancing our understanding of the biological effects of ultrasound-based ablative therapy and inspiring clinically relevant applications in the future.

**Key Points:**

1) Development of an integrated platform for real-time, on-the-fly imaging of FUS-induced cell ablation and response processes at cellular and subcellular levels
2) Focused ultrasound induces cellular mechanolysis through previously unappreciated subcellular cavitation bubbles that grow under confined contact with cells
3) Post-ablation phantom cell layer could serve as a guiding cue for collective wound healing after ultrasound-inflicted ablation

## 1. Introduction

In recent years, focused ultrasound (FUS) has attracted extensive attention as a non-invasive therapeutic modality and rapidly advanced in various research areas.^[1–3]^ Currently, the most common application of therapeutic FUS is in the ablation of tumors.^[4,5]^ Furthermore, FUS exhibits deep penetration and high spatial precision in comparison to other therapeutic physical stimuli. Low-frequency focused ultrasound (frequency < 1 MHz) could non-invasively focus within the cranial region and work in combination with other therapeutic modalities such as microbubbles and phase-change liquid droplets.^[6,7]^ y promoting transcellular endocytosis, inducing endothelial cell membrane opening, forming cavities or channels within the cytoplasm, or opening tight junctions between endothelial cells, FUS facilitates the delivery of therapeutic agents, *e.g.*, cisplatin, doxorubicin, mistletoe lectin-1 (ML1), trastuzumab, and genetic materials across the blood-brain barrier (BBB) or blood-tumor barrier (BTB).^[8–18]^ FUS could also impose inhibitory or activating effect on neural activity with potential applications in analgesia.^[19–22]^ FUS stimulation of the hepatic portal venous plexus holds promise as a non-pharmacological treatment for glucose homeostasis dysregulation.^[23]^ Moreover, FUS has been applied in thrombolysis and therapeutic research related to inflammation.^[24–26]^ Overall, FUS-based therapeutic techniques offer vast avenues for clinically relevant applications.

The therapeutic principle of FUS technology is based on the energy input generated by ultrasound waves focused at specific areas in tissues, leading to thermal or mechanical effects, followed by a series of biological effects. The thermal effect of FUS is directly caused by temperature-dependent responses such as local hyperthermia and coagulative necrosis.^[27–29]^ The mechanical effects of FUS primarily include cavitation process such as stable or inertial cavitation. Stable cavitation refers to the stable oscillation of microbubbles in the focal region under the effect of pressure waves. Inertial cavitation involves the drastic volume changes of microbubbles under the influence of strong pressure waves, ultimately leading to their rupture and generating shock waves.^[30]^ Despite considerable research on the physical process of FUS, the investigation into the biological effects such as cell ablation induced by HIFU, and the subsequent wound healing response, remains limited. This hinders a comprehensive understanding of the mechanisms underlying ultrasound-inflicted ablative therapy and limits the advances in ultrasound-based therapeutic technology. Therefore, it is critically needed to explore the mechanism of HIFU-induced cell ablation and associated wound healing responses.

To investigate the mechanisms of ultrasound therapy, *e.g.*, HIFU-induced cell ablation, it is essential to develop experimental models that can recapitulate multicellular damage and repair processes under ultrasound treatment in vitro. Traditionally, in vitro studies on the biological effects of ultrasound typically expose cells to FUS treatment before observing them under a microscope.^[31–33]^ Such two-step operations are performed sequentially on separate platforms and thus lacks real-time, on-the-fly observation of the immediate / short-term responses (0-10 min) during FUS treatment. Currently, devices for studying ultrasound-induced responses mainly rely on either an upright or an inverted microscope. For example, Zhihai *et al*. integrated a FUS device with an upright microscope, placing the FUS transducer beneath the culture dish at 45-deg, which allows incident sound waves to reach the culture dish through a waveguide material, while ell imaging was performed by immersing the objective into the upright-placed cell culture dish.^[34]^ Recently, Yao-Shen *et al*. combined a FUS device with an inverted microscope, with the FUS transducer immersed vertically into the cell culture medium.^[35]^ However, these setups required immersing the objective lens or ultrasound transducer into an open cell culture environment, leading to a significant risk of contamination. Despite that they might be usable for short-term experiments, they are insufficient to support continuous studies on the cell ablation mechanisms and subsequent wound healing responses induced by FUS. This technology gap hinders the exploration and understanding of ultrasound-inflict cell ablation mechanisms. Therefore, it is imperative to develop new technologies that integrate non-invasive ultrasound stimulation and real-time, on-the-fly live-cell imaging and long-term cell culture and tracking into a single platform.

Herein, this study designed and constructed a fully enclosed cell culture device (ECCD) that enables non-invasive ultrasound stimulation and real-time, on-the-fly live-cell imaging of FUS-induced cell ablation and response processes at cellular and subcellular levels. Unlike previously reported open culture systems, this platform allows for real-time and programmable FUS stimulation of cells in the targeted focal region, while also enabling online monitoring of cellular responses for long term. We applied this platform to obtain a stereotypic focal-area geometry under FUS, and analyze the biophysical effects involved in different sub-regions of the focal-area, including short and long-term cellular responses during and after HIFU-induced cell ablation. This study not only provides direct experimental evidence and insights into classical thermal and mechanical effects of FUS, but also reveals previously unappreciated biomechanical mechanisms underlying the cell damage and collectively wound healing involved in FUS-based ablative therapy.

## 2. Results

### 2.1. Design of an Enclosed Cell Culture Device Compatible with FUS Stimulation and Long-Term Live-Cell Imaging

To simultaneously achieve non-invasive FUS stimulation and real-time, on-the-fly live-cell imaging for long term, we developed an enclosed cell culture device (ECCD). Through optimized design of the acoustic path, acoustic windows, and sound-absorbing materials, this device enables FUS to act on a focal domain within cultured cells (**Figure 1A**). Meanwhile, real-time imaging captures morphological changes and biological responses of cells within the focal domain as well as those outside of it. Specifically, this design features a clear, uninterrupted acoustic path, which allows the FUS emitted from the transducer to travel through a coupling device, pass through the entrance acoustic window (EnAW), propagate and focus at the cell culture region at the bottom of the ECCD. The incident ultrasound cone forms a 45-deg inclined angle and thus is also reflected at 45-deg before passing through the exit acoustic window (ExAW) and absorbed by a sound-absorbing material attached to the exterior of ExAW, thereby avoiding additional reflections that may affect the focused acoustic field within the cell culture device.

**Figure 1.**
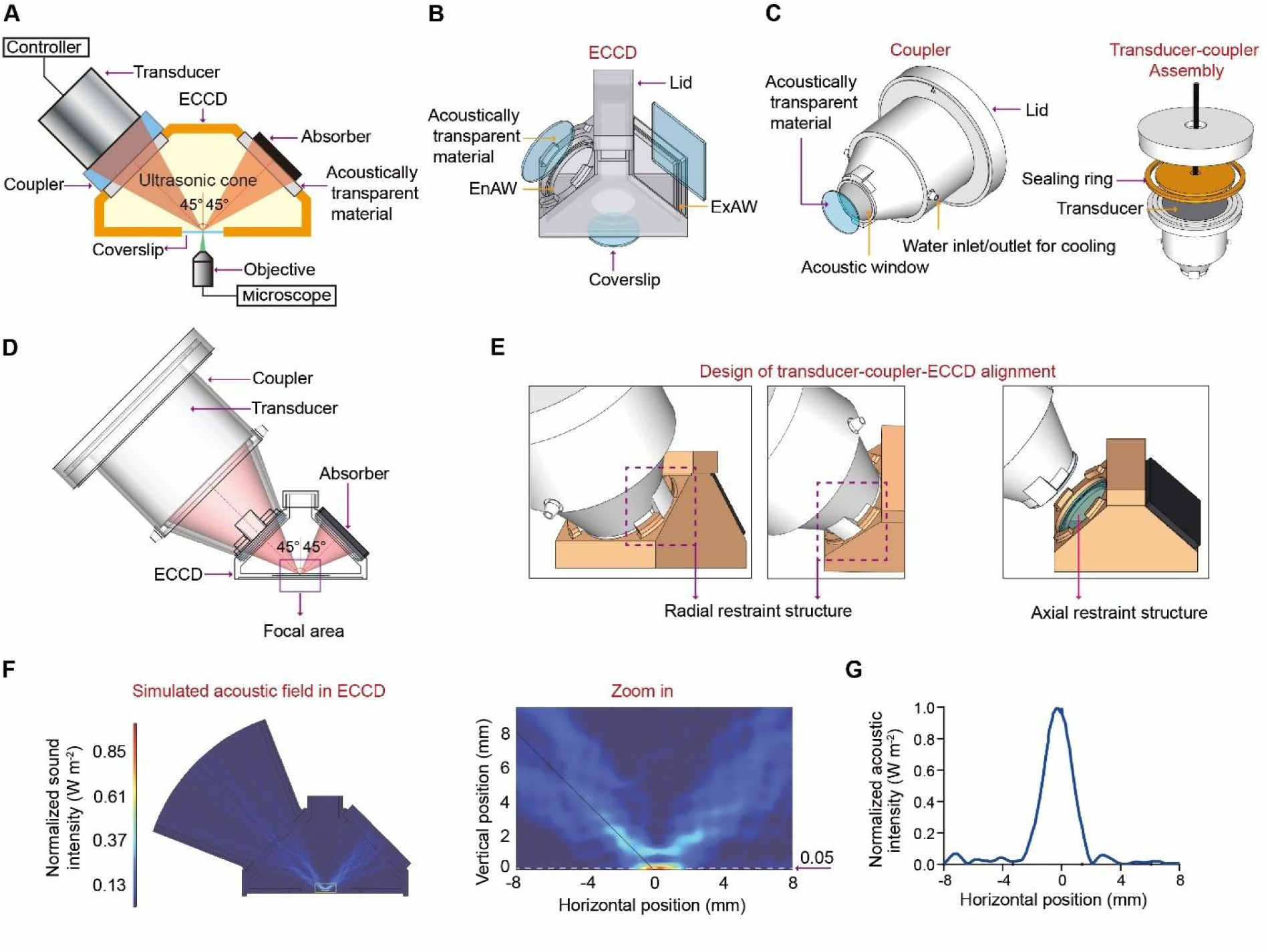
Design of the enclosed cell culture device (ECCD) compatible with focused ultrasound (FUS) stimulation and real-time, on-the-fly imaging. **(A)** Schematic of the ECCD design and the acoustic pathway of FUS stimulation. **(B)** Structural components of the ECCD. **(C)** Structural components of the FUS transducer coupler. **(D)** Schematic showing the aligned assembly of the FUS transducer and ECCD with the acoustic pathway and expected focal area. **(E)** Design of restraint structures for co-axial transducer-coupler-ECCD alignment. **(F)** Simulation results of the acoustic field elicited by FUS in the ECCD. Zoom-in image displays the acoustic intensity field in the focal region. The acoustic intensity along the white dashed line (at the height of 0.05 mm above the bottom of ECCD) was extracted and plotted in (G). **(G)** Line-profile of acoustic intensity along the white dashed line shown in (F), showing the focused deployment of acoustic energy in the expected focal region.

The ECCD is assembled from multiple components. To meet requirements for FUS stimulation and standard cell culture, we adopted a four-sided pyramid structure (**Figure 1B**). The circular EnAW and rectangular ExAW on the left and right sides of the device, respectively, are inclined at 45-deg and sealed with 1 mm thick sheet of acoustically transparent material. This allows the FUS beam to transmit with extremely low loss along the expected acoustic path while avoiding direct contact between the ultrasound transducer and the liquid inside the culture device. A circular rim is machined to extend from the inner edge of the EnAW and ExAW for restraining the position of acoustically transparent sheets. To ensure correct positioning and stable contact between the ultrasound transducer coupler and the EnAW, a set of protruding structures were manufactured at the inner rim of the EnAW to restrain the movement of the coupler along both radial and axial directions. A circular opening and recess structure is made at the bottom of the ECCD to allow the attachment of a coverslip for cell culture (Figure 1B).

Further, a coupler is designed to integrate the FUS transducer with the ECCD. To match the shape of the ultrasound transducer and the ultrasound cone, the coupler contains a conical headpiece (**Figure 1C**), which has a circular opening covered with a 1 mm thick acoustically transparent sheet. The coupler also features a set of protruding structures complementary to those on the rim of the EnAW of ECCD, ensuring the co-axial assembly and tight fit between them (**Figure 1D&E**). During operation, the coupler is filled with degassed water and connected to a circulatory cooling system. The FUS transducer is mounted to the rear part of the coupler, before being sealed by a water-tight lid and a silicone ring.

To examine whether the focused acoustic field in the device meets the expectation from our design, we conducted multiphysics simulations using COMSOL. For simplicity, a two-dimensional model was used (**Figure S1A-D**). Specifically, the computed sound pressure field, sound pressure level field (**Figure S1E&F**), and sound intensity field (**Figure 1F**) in our multiphysics simulation model were normalized and plotted as heat maps. These results showed that the ultrasound waves converged along the expected acoustic cone to reach the focal domain, reflected at the expected 45-deg inclined angle, before traveling towards the ExAW along the designed path and absorbed by the sound-absorbing sheet (modeled as a perfect matching layer in the simulation). These results clearly indicate a focal ultrasound stimulation at the target region of the cell culture area, with significantly lower level of ultrasound outside of the focal region (**Figure 1G**). Similar results are also observed in the computed sound intensity field as well. Such focused stimulation performance meets the goal of our device design. To evaluate the importance of sound-absorbing materials in above design, we attempted to replace it with a sound-reflective boundary in the simulation (**Figure S1G&H**). Indeed, the absence of sound-absorbing materials at the ExAW resulted in highly disordered, non-focal sound pressure field and sound pressure level field in the device (**Figure S1I&J**). Under such condition, the incident acoustic energy could not be accumulated at the focal, and a stable energy distribution cannot be calculated in the steady-state sound intensity field. Therefore, the sound-absorbing material in the design of ECCD is crucial for ensuring a clear, undisrupted, and predictable acoustic path of FUS. In addition, we compared the acoustic field of ECCD with that generated in a conventional upright imaging system, which was previously used to stimulate cells with ultrasound (**Figure S2A&B**). Our simulation results showed that in contrast to the focal acoustic field formed in the ECCD, the conventional system only generated disordered acoustic field despite the use of a focused ultrasound transducer (**Figure S2C&D**), due to the lack of design to optimize the acoustic path. Therefore, our results implicate the importance of ECCD device design to assure desirable FUS-based focal stimulation and unobstructed path for both acoustics and live-imaging at the same time.

### 2.2. An Integrated Platform for FUS Stimulation and Live-Cell Imaging Based on ECCD

In order to integrate ECCD with laser confocal microscopy, a two-level carrier system was designed. This system allows the transducer, coupler, and ECCD to move freely in three dimensions within the microscope’s field of view, facilitating optical focusing and field-of-view selection. To this end, a three-axis displacement platform, referred to as the primary displacement system, was employed (**Figure 2A&B**). This system also mechanically isolates the weight of the ultrasound transducer, coupler, and culture device, from the microscope platform, thereby ensuring minimal disturbance to the optical systems as well as universal applicability to different kinds of optical setups. Furthermore, ensuring alignment and close contact between the ultrasound transducer coupler and the culture device is crucial for maintaining the stability and predictability of the acoustic pathway during experiments. Therefore, a secondary displacement system was mounted on the primary platform to control the relative motion between the coupler and the culture device. This system, with only one degree of freedom in the vertical direction (*Z*-axis), independently controls the vertical motion and applies additional load to achieve precise alignment between the coupler and the EnAW. With such a two-level carrier system, independent adjustments could be made to the acoustic pathway as well as the optical path relative to the microscope, decoupling the control of ultrasound stimulation and that of microscopic imaging.

**Figure 2.**
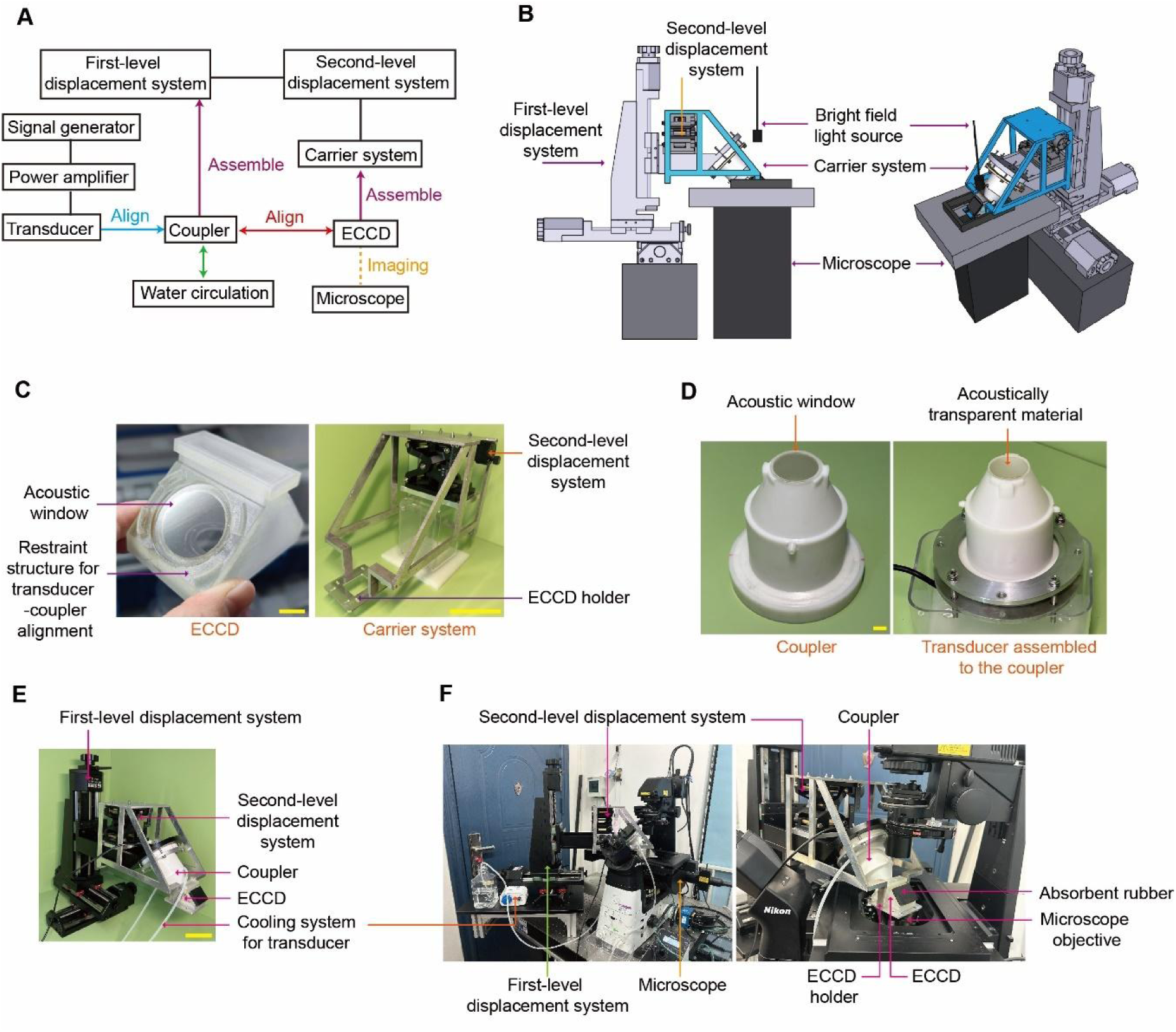
An integrated platform for FUS stimulation and live-cell imaging based on the ECCD. **(A)** Flowchart showing the experimental setup and its operational procedure. **(B)** Schematic showing the assembly of the integrated platform, including the first-level (tri-axial) and second-level (uni-directional) displacement control systems that are designed to control the 3D movement of the ECCD within the light path of the microscope and the alignment of the transducer to the ECCD, respectively. **(C)** Photographs showing the ECCD and its carrier system. The ECCD holder is connected to the seond-level displacement system. Scale bars: 10 mm (left), 10 cm (right). **(D)** Photographs of the coupler before (left) and after (right) assembled to the FUS transducer. Scale bar: 10 mm. **(E)** Photograph showing the assembly of the first and second-level displacement systems, carrier system, FUS transducer and coupler, cooling system for the transducer, and the ECCD. Scale bar: 10 cm. **(F)** Photographs showing a fully assembled, integrated platform for FUS stimulation and real-time microscopic imaging based on the ECCD.

High-precision 3D printing was chosen to manufacture the ECCD and the acoustic coupler, respectively. The ECCD was made from MED610 photopolymer resin, which is commonly used in orthopedic and oral surgery and expected to exhibit good biocompatibility, thermal stability, and chemical stability under cell culture and ultrasound stimulation conditions.^[36]^ It also demonstrates mechanical stability to prevent deformation, breakage, fatigue, and creep during assembly and experiments. The coupler was made from ABS photopolymer resin (acrylonitrile-butadiene-styrene copolymer) using high-precision 3D printing (**Figure 2C&D**), providing endurable performance based on the robustness, density, and stability of the resin material.

For the acoustic windows on the ECCD and the coupler, 1 mm thick sheets of acoustically transparent materials were used to both ensure seamless contact and avoid mechanical bending due to liquid pressure under working conditions. Of note, these acoustic windows should be made from materials that do not significantly reflect sound waves at their interface with the culture medium (with acoustic property similar to that of water). To select such acoustically transparent materials, we first screened several common materials for their acoustic impedance, from which polydimethylsiloxane (PDMS) was found to feature an impedance value close to that of water (**Table S1**). Towards further optimization, we screen PDMS materials with different base : curing-agent mixing ratio, and found that the acoustic impedance of PDMS sheets made of at 4:1 mixing ratio is closest to that of water at the same working temperature (**Table S2**). Therefore, PDMS with base: curing agent ratios at 4:1 was used throughout this study for manufacturing acoustic windows. Before conducting experiments, the primary displacement platform carrying the FUS stimulation system and the secondary displacement platform carrying the ECCD (**Figure 2E**) are assembled, connected to circulatory cooling system for the transducer, and mounted on the optical table and aligned with the optical path of the microscope (**Figure 2F**).

In addition, we calibrated the acoustic field generated by the FUS transducer used in this study. In particular, the calibration was conducted using a hydrophone to measure the sound intensity field near the focal plane of the transducer. The results showed that when the input voltage, *U*, to the transducer was 20 V, at the frequency of 1 MHz, the repetition frequency of 10 kHz, and the duty cycle of 30% for burst waves, the −3 dB focal region (where the sound intensity dropped to 50% of the maximum intensity) on the *x*-*y* plane (perpendicular to the axis of the focusing acoustic cone) was approximately a circular area with a diameter of 3 mm; while on the *x*-*z* plane, the −3 dB focal region was an elliptical shape with a major axis of about 18 mm and a minor axis of about 3 mm (**Figure S3A**). Given that thermal effect due to HIFU is inevitable during experiments, we studied the dynamics of average temperature change at the focal region. Specifically, we used a high-sensitivity thermistor measurement method. Specifically, a T25 culture flask was attached with a thermistor at the bottom and filled with water, before being placed on the path of the FUS (**Figure S3B**). To ensure high thermal sensitivity and minimal scattering of the acoustic waves, the thermistor was made from a thin platinum wire (50 μm in diameter). To measure the average temperature over the focal region, the platinum wire was shaped into a serpentine configuration and attached to a 3×3 mm^2^ square area at the bottom of the flask, which matches the size of the focal region of the FUS transducer (**Figure S3C-F**). Our results showed that under FUS stimulation, the temperature at the focal region could rise and reach a steady state within ∼ 5 s after the FUS is on, and drop to the environmental temperature within 3 s after the FUS was stopped, thus demonstrating the rapid temperature response at the focal region. Moreover, the target temperature of the focal region could also be modulated by varying the ultrasound intensity (via transducer voltage *U*) and the duty cycle of burst waves (**Figure S3G&H**).

### 2.3. FUS-Induced Cell Ablation Does Not Show Significant Correlation with Thermal Effect

To mimic the interaction between FUS and tissues (especially epithelial tissues, which are commonly involved in ultrasound therapy), this study selected epithelial monolayer of Madin-Daby canine kidney cells (MDCK cells) as an in vitro model. Firstly, to validate the compatibility of ECCD with cell culture, MDCK cells were plated in ECCD and regular 60 mm culture dish and cultured for 2 days. No significant difference was observed between above two groups in cell proliferation and epithelial colony morphology at 50% and 100% confluence, respectively (**Figure 3A**). In order to observe the shape and size of focal region produced by FUS on MDCK monolayer in ECCD, we next set to simultaneously investigate the temperature change of focal region and cell ablation under FUS stimulation. Although abovementioned results from thermistor experiment showed the dynamics of average temperature change in the focal region, the thermistor method was not suitable for measuring the temperature field distribution of the focal area. To characterize the temperature field, custom-made thermochromic tapes were used. These tapes contained thermochromic inks with pre-calibrated color-changing temperatures of 31, 38, 45, and 65 °C in red, yellow, green, and purple, respectively, and were mixed at volume ratio of 4:2:2:1. Upon heating, these inks change color from brick red at room temperature to green, blue, purple, and milky white at 34, 38, 43, and 62 °C, respectively (**Figure 3B**). These thermochromic tapes were applied to the bottom of the ECCD before FUS experiments, and the color changes were recorded to reconstruct the temperature field changes during ultrasound stimulation. Additionally, to detect cell ablation after ultrasound stimulation, Live dye was used to label the entire monolayer of MDCK cells at 100% confluence before the experiment. Of note, while Live dye emits a strong green fluorescence (FITC) inside cells with intact plasma membrane, it produces little or no fluorescence signal outside the cell or once cell membrane was ruptured, thereby serving as an indicator of cell ablation. It is also important to note that the disappearance of FITC signal only reflect cell ablation, not necessarily cell detachment.

**Figure 3.**
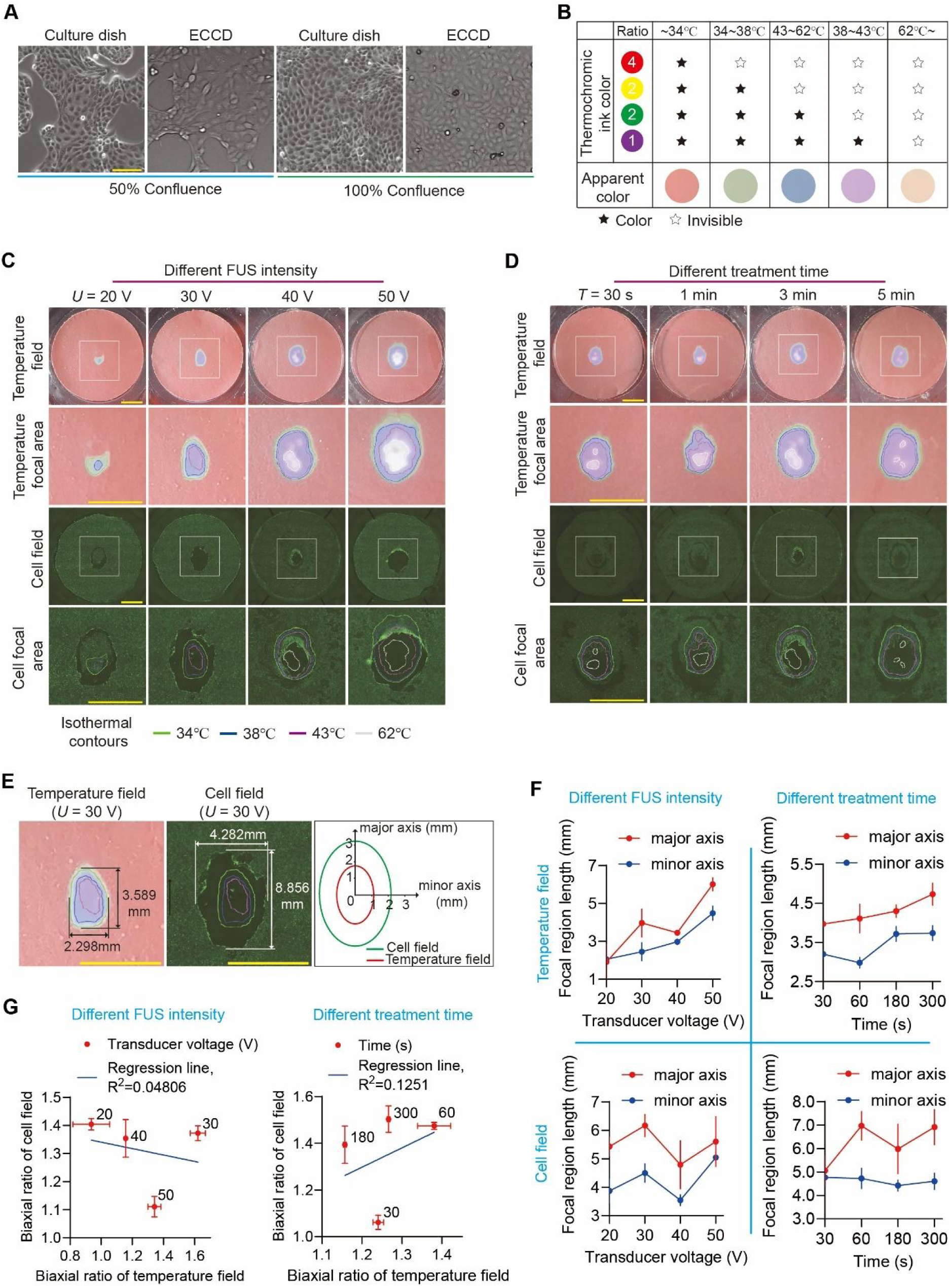
FUS-induced cell ablation does not show significant correlation with ultrasound-caused thermal effect. **(A)** Bright-field images of MDCK cells cultured in regular 60 mm culture dishes and ECCD at confluence of 50% and 100%, respectively. Scale bar: 100 μm. **(B)** Table showing the temperature-dependent coloration of the four-color thermochromic ink. The mixing ratio of mono-chromic ink is red: yellow: green: purple = 4:2:2:1. **(C)** Representative images showing the distribution of temperature and cell ablation, respectively, in the ECCD upon FUS treatment at different transducer voltage (*U*) while the frequency (1 MHz, continuous waves) and treatment duration (*T* = 3 min) remain constant. The gradient coloration of the thermosensitive tape was used to plot the isotherm contours of the temperature field. Live dye (green) was used to visualize living cells and the corresponding FITC fluorescence confocal micrographs (cell field) were used to analyze the cell ablation within a MDCK monolayer. Cell ablation area is reflected by the loss of fluorescence after FUS treatment. Square boxes indicated the region shown in zoom-in images. Similar results were observed in *n* = 3 independent experiments. Scale bar: 5 mm. **(D)** Representative images showing the distribution of temperature and cell ablation, respectively, in the ECCD upon FUS treatment for different duration (*T*) while the frequency (1 MHz, continuous waves) and transducer voltage (*U* = 40 V) remain constant. Similar results were observed in *n* = 3 independent experiments. Scale bar: 5 mm. **(E)** Representative distributions of the temperature field (left) and cell field (right) at the focal domain of the ECCD upon FUS treatment at *U* = 30 V and *T* = 3 min. The contours of the temperature field and cell field were demarcated as indicated, respectively, with the dimensions of their major axis and minor axis labeled accordingly. **(F)** Plots showing the change of the dimensions of the focal region of the temperature field (upper panels) and cell field (lower panels), respectively, under indicated conditions of transducer voltage (left) or treatment duration (right). Data were plotted as mean ± 95% confidence interval. *n* = 3 independent experiments. **(G)** Plots showing the relationship between the aspect ratio of the focal region of the temperature field and that of the cell field, under indicated conditions of transducer voltage (left) or treatment duration (right). Linear fitting was applied using the least squares method. Data were plotted as mean ± s.e.m. *n* = 3 independent experiments.

Given that HIFU-based thermal effects are widely believed to be associated with ultrasound-induced cell ablation in previous studies^[4,27]^, we next compared the temperature changes within the focal region to the spatial distribution of cell ablation. The “temperature field” was henceforth defined based on the images of the thermochromic tape, with particular attention to the region where color change (*i.e.*, temperature change) occurred. This temperature field was plotted as isotherms and clearly visualizes the focal area under FUS treatment. Correspondingly, the images of live cells featuring FITC fluorescence were henceforth defined as the “cell field” to analyze the change in live cell distribution after treatment. Specifically, we define areas where cells were ablated (shown by the disappearance of fluorescence during FUS treatment) as the “cell ablation area”, and an elliptical-like enveloping area that encloses all cell ablation area was defined as the focal area of cell field. Our results indicated that under FUS treatment, the focal area of the temperature field reached a steady state within a short time (1-10 s). After the termination of FUS, the focal area of the temperature field returned to ambient temperature within a relatively short time (**Figure S4A**). This was consistent with the temperature change dynamics measured by thermistor-based method (Figure S3F-H).

Of note, our results showed that the contours of the temperature field and the cell field were approximately an elliptical shape, with the major axis oriented along the direction of incident ultrasound waves (**Figure 3C&D**). By measuring the lengths of the major and minor axes of such elliptical contours (**Figure 3E**), we observed a positive correlation between FUS intensity and the size of focal region of the temperature field, as well as that of the cell field. As expected, similar correlation was also observed between the size of abovementioned focal regions and FUS treatment time (**Figure 3F**).

However, by comparing the cell ablation area with the focal area of the temperature field, it is clear that the focal area of the temperature field showed continuously decreasing temperature from the central to peripheral domains, while the distribution of the cell ablation area was not necessarily continuous or following such pattern of temperature distribution. In contrast, the cell ablation area features a hollow elliptical shape under high ultrasound intensity (*U* = 50 V, treatment time *t* = 3 min), while it became a circular elliptical band under lower ultrasound energy (*U* = 20 V, *t* = 3 min), with non-uniform cell ablation domains observed under intermediate range (*U* = 40 V, *t* = 3 min). Further quantitative analysis revealed no significant linear correlation in terms of shape and size between the cell ablation area and the focal region of the temperature field (**Figure 3G**). These observations suggeste that although changes in the temperature field occurred as expected within the focal region, the continuous changes in the temperature field and the non-continuous, non-uniform changes in the cell ablation area were inconsistent with the conventional understanding of ultrasound-based thermal ablation theory. This implies that mechanisms other than thermal effects might exist for cell ablation under FUS.

To further demonstrate that the thermal effects within the FUS focal area were insufficient to ablate cells within this region in the short experimental duration of this study, we placed the ECCD (with a monolayer of MDCK cells at over 90% confluence) under heat baths at 65 and 85 °C, respectively, for 5 min. Compared with the untreated group, the level of cell ablation (reflected by the relative cell ablation area) caused by short-term heat treatment was less than 10% in both 65 °C and 85 °C groups. This is significantly different from the proportion of cell ablation area within the focal region under FUS treatment (**Figure S4B-D**). These results indicate that cell ablation produced under FUS stimulation in a short time and the associated thermal effects in the focal area were not significantly correlated in a global manner. Therefore, it is necessary to explore cell ablation mechanisms beyond thermal effects.

### 2.4. Ultrasound-Induced Cell Ablation Shows Multimodal Association with Cavitation Bubbles

To explore FUS-induced cell ablation mechanisms beyond global thermal effects, we performed experiments using MDCK cells expressing red fluorescent protein in cell nuclei (H2B-RFP). Firstly, we applied Live dye to visualize the region of live cells as well as that of cell ablation. **Figure 4A** shows a typical topology of the cell-ablation area after HIFU treatment, which features an elliptical ablation band that envelop a central area containing streak-like ablation domains and a small, high-temperature domain. Specifically, when the FUS was activated for a few seconds, the streak-like ablation pattern appears near the nucleation-like high-temperature domain, and expands from the side proximal to the incident FUS waves, as shown in **Figure S5**. Of note, there were no cavitation bubbles in the streak-like ablation domains during FUS treatment, suggesting it likely initiates from a combination of high-temperature and shear force and then continued to expand due to acoustic streaming driven by the nonlinear effect of ultrasound.^[37]^ In the meantime, after the FUS was activated for a few seconds, a large number of cavitation bubbles gradually appeared near the side proximal to the incident FUS waves, causing cell ablation at this site, which evolved into a crescent-like shape and eventually a complete elliptical ablation band. Even after the cessation of FUS, some bubbles remained in the crescent-like region (**Figure 4B**). These bubbles are opaque under the bright field, but almost transparent in the fluorescent channel, with only a halo around the edge due to environmental light scattering effect at the air-liquid interface around the bubble.^[38,39]^ These data suggest that above-observed cell ablation is closely associated with cavitation bubble formation under FUS.

**Figure 4.**
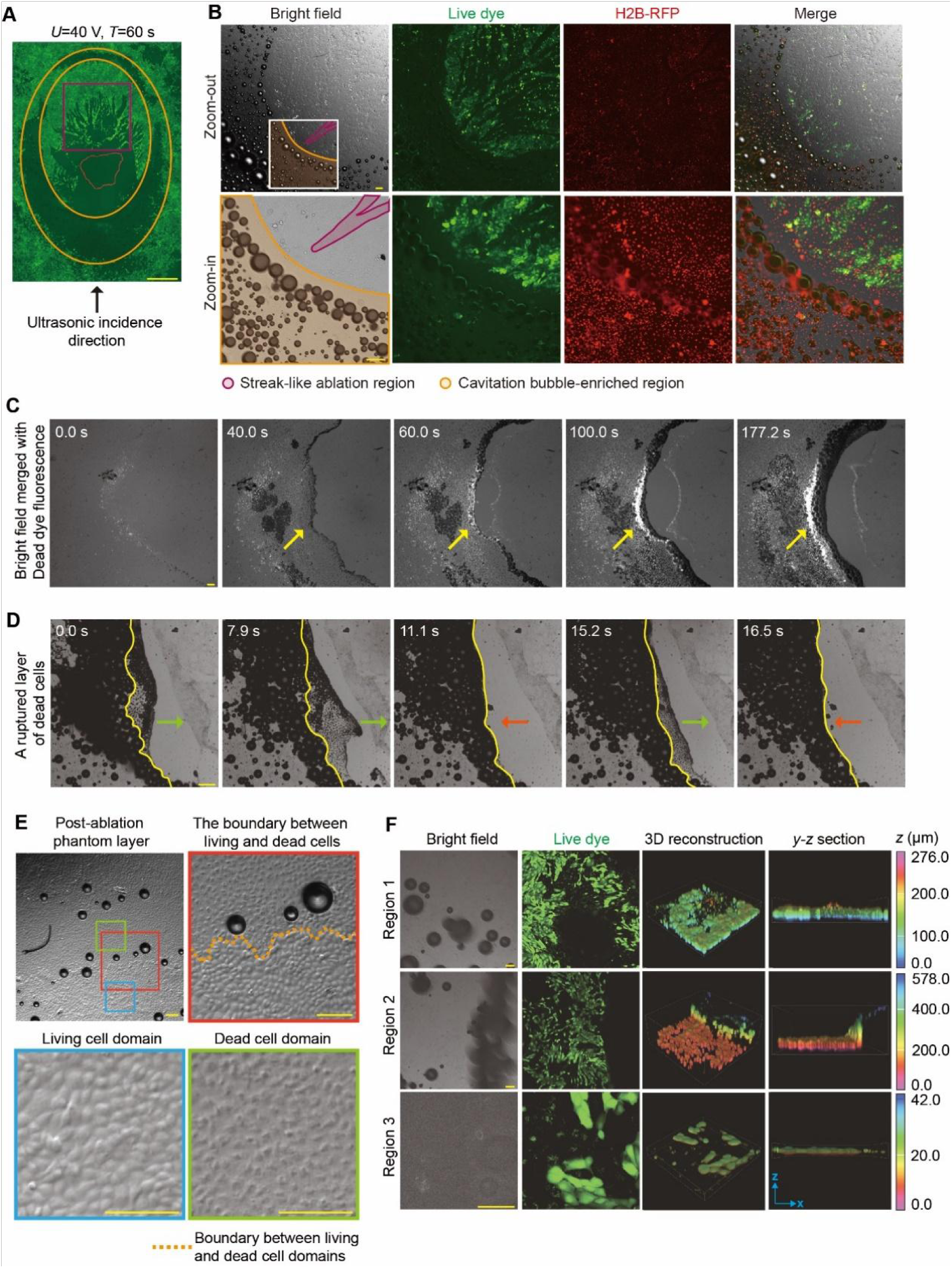
Significant association between cell ablation zone and bubble generation zone under FUS. **(A)** Confocal micrographs showing the distribution of living cells (tracked by FITC fluorescence) after FUS treatment (*U* = 40 V, *T* = 60 s, 1 MHz continuous waves). The black arrow indicates the direction of incident ultrasound waves. Live dye is used to label living cells, and non-fluorescent regions reflect the cell ablation zone. Scale bar: 1 mm. The purple box marks the streak-like ablation domains in the focal area. The area between the two orange ellipses is the ablation region where cavitation bubbles appear. The area enclosed by the red irregular lines is the area with a temperature greater than 62 ℃ during FUS treatment. **(B)** Confocal fluorescence micrographs showing the distributions of cavitation bubbles, living cells, and cell nuclei, within the focal region after FUS treatment (*U* = 40 V, *T* = 3 min, 1 MHz continuous waves). Bubbles are opaque and spherical under bright field. Live dye (green) is used to detect living cell domain and non-fluorescent cell ablation domain. H2B-RFP MDCK cells are used to track the cell nuclei (red). Similar results were observed in *n* = 5 independent experiments. Scale bar: 100 μm. **(C)** Time-lapse micrographs showing the progressive cell ablation marked by Dead dye fluorescence within the focal domain of a monolayer of MDCK cells under FUS treatment (*U* = 40 V, continuous waves at 1 MHz). Yellow arrows indicate bands of bubbles. The time-stamp represents the time lapsed from the first image. Scale bar: 100 μm. **(D)** Time-lapse micrographs showing the progressive rupture of the MDCK monolayer under continued FUS stimulation (*U* = 40 V, 1 MHz continuous waves). Yellow lines represent the boundary of the bubble-enriched region. Red and green arrows indicate the expansion and retreat movement of bubble clusters. The time-stamp represents the time lapsed from the first image. Scale bar: 100 μm. **(E)** Representative images showing the post-ablation phantom layer (PAPL) composed of FUS-ablated dead cells and its neighboring living cell domain (*U* = 40 V, *T* = 3 min, 1 MHz continuous waves). Colored square boxes mark the living cell domain (blue), dead cell domain (green), and the boundary between living and dead cell domains (yellow dashed lines in red box) in zoom-in images. Scale bar: 100 μm. **(F)** Confocal fluorescence micrographs showing the post-ablation MDCK monolayer (*U* = 40 V, *T* = 3 min, 1 MHz continuous waves). Living cells are marked by live dye (green). 3D reconstructed images of the PAPL and its adjacent living cell monolayer are also shown. Region 1 shows the PAPL that remained in its original place; region 2 shows PAPL ruptured and peeled off the substrate by bubbles, along with some live cell monolayer connected to it; region 3 shows the PAPL forming a “dome-like” shape, with its central domain detached but peripheral domain connected to the live cell monolayer on the substrate. Color bar represents the *z*-position within the 3D reconstructed image. Scale bar: 100 μm.

To further investigate the relationship between FUS-induced cavitation bubbles and cell ablation, we used Dead dye to monitor cells that died during FUS treatment in real time. Of note, when the FUS in on, a band of bubbles first appeared (**Figure 4C&D**), wherein bubbles were constantly generated, expanded and finally burst under FUS. Due such mechanical effect within this band of bubbles, cells in this region were ablated rapidly, as reflected by the presence of the Dead dye signal. Such ablated cell monolayer eventually broke down and ruptured, leaving a narrower void band. Meanwhile, we did not observe notable temperature rise in this region, ruling out that the bubbles in this region are caused by the heating and boiling of the liquid in this region. Therefore, we reason that this phenomenon could be attributed to the inertial cavitation phenomenon.

Strikingly, we observed a large number of steadily expanding bubbles in the crescent-like region proximal to the incident FUS waves. These bubbles, in contrast to above-mentioned inertial cavitation, remain stable and growing without bursting, in the absence of significant temperature increase. This suggests that these cavitation bubbles and corresponding cell ablation in this region might reflect a mechanism different from canonical effects such as thermal-ablation or cavitation bubble collapse.

To further illustrate the relationship between stable FUS-induced bubbles and cell ablation observed in the proximal domain of the crescent-like region, we treated cells with pulsed FUS of varying intensity with a duty cycle of 10% and a pulse repetition rate of 1 kHz (**Figure S6**). Compared with abovementioned continuous FUS waves, pulsed FUS with the same transducer voltage showed greatly reduced temperature within the focal region. Under pulsed FUS, the streak-like ablation domains in the central region of the focal plane are still evident, further indicating it is likely due to non-cavitation mechanical effect such as shear stream instead of thermal effect. Importantly, with pulsed FUS, there was no notable formation of stable bubbles, nor formation of elliptical ablation band with stable bubbles or cell ablation associated with it. These results indicate that FUS-induced cell ablation in the unique crescent-like domain requires the formation of stable cavitation bubbles.

Notably, a layer of dead cells remained attached after cell ablation within the elliptical ablation band (**Figure 4E**). Bright-field imaging results showed that while this layer of dead cells still appear to connect with the rest of the cell monolayer, its morphology differed significantly from live cells. This suggests that cell ablation under ultrasound may not result in the complete disappearance of cells but leaves behind a post-ablation phantom layer (PAPL) composed of cellular remnants. Importantly, the connections formed by the cytoskeleton in the phantom layer appear intact, maintaining connections to adjacent live cell monolayer. Although such connections might hold the phantom layer in its original place, it was also observed that after long-term FUS treatment, or at the location where large bubble clusters form, the phantom layer could break away from the substrate (**Figure 4F**). In this process, even part of the live cell monolayer connected to the phantom layer could be peeled off the substrate. These results indicate a significant relationship between FUS-induced cavitation bubbles and the cell ablation potentially via mechanical interactions between them. This suggests that it is critical to investigate deeper into the mechanical interactions between FUS-induced bubbles and cells. Additionally, the formation of the PAPL in above FUS-induce tissue damage model might represent a previously unappreciated opportunity to recapitulate the wound created by HIFU-based cell ablation.

### 2.5. Cellular Mechanolysis Induced by Subcellular Cavitation Bubbles

Above experiments provided macroscopic level evidence regarding a mechanical mechanism underlying HIFU-induced cell ablation mediated by cavitation bubbles. To further explore the relation between bubble formation and cell ablation in the elliptical ablation band at the microscopic level, we observed cellular responses during FUS treatment at single-cell resolution. Live dye was used to track cell viability in real-time. Our results showed that, as ultrasound stimulation proceeded, a subcellular cavitation bubble suddenly appeared where cells attached. While the bubble kept growing, the Live dye fluorescence within the cell that associated with the bubble decreases until almost no fluorescence signal could be detected in the cell (**Figure 5A**). Quantitative analysis of the fluorescence intensity revealed a rapid decay within a few seconds after the appearance of the bubble, followed by a minimally detectable plateau (**Figure 5B**). These results suggest that as the bubble grows, the cell membrane was disrupted, leading to the outflow of cellular content and eventually cell death. Statistical analysis indicated that the fluorescence signal of the cell decreased from the level of live cells to the background level in an average time of about 10 s after the formation of the bubble (**Figure 5C**). Compared to classical processes such as apoptosis or necrosis, this is a remarkably rapid process, implying a different mechanism for cell ablation through mechanical interaction between cells and FUS-induced subcellular cavitation bubbles.

**Figure 5.**
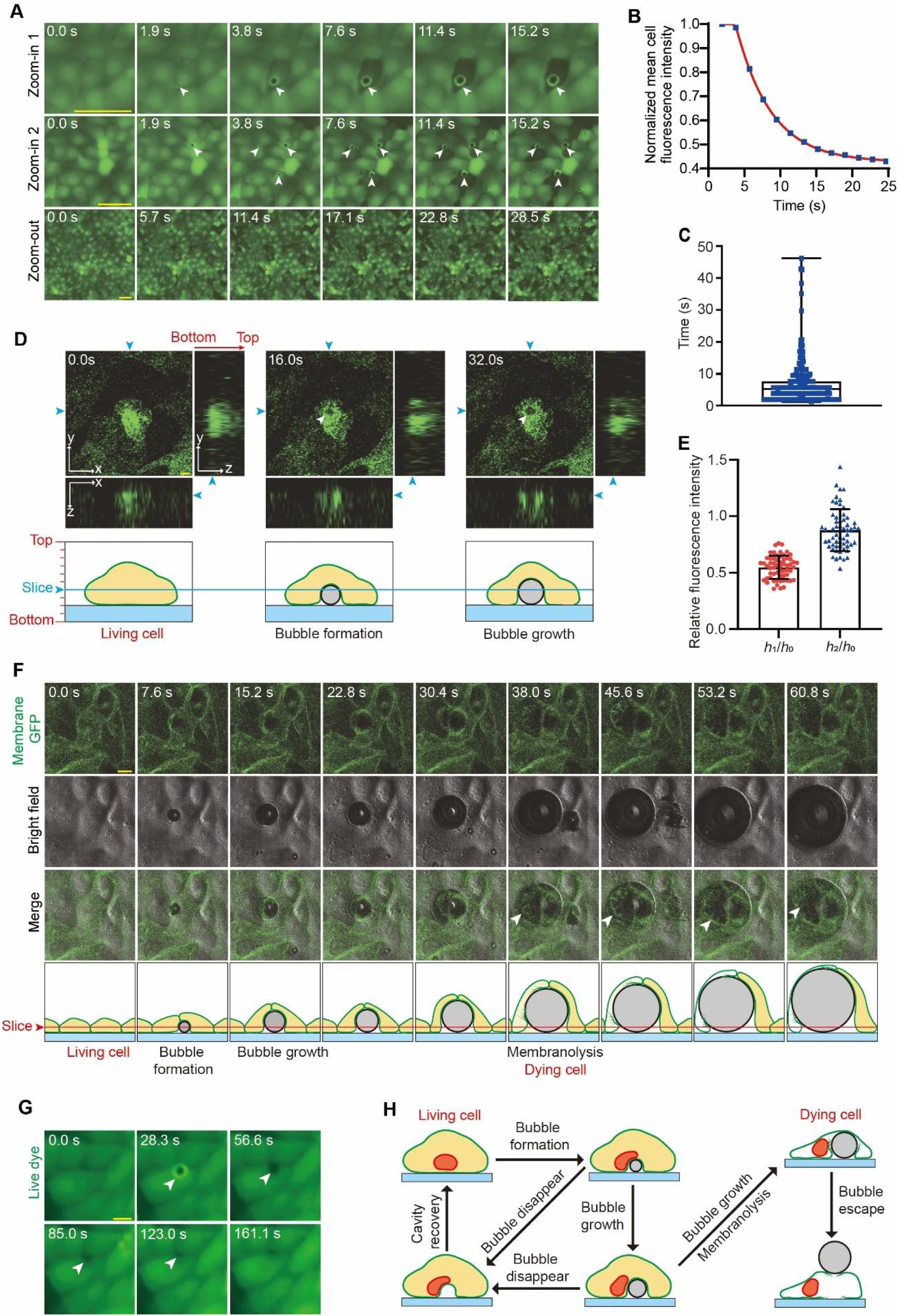
Subcellular cavitation bubble formation and FUS-induced mechanolysis of the cell. **(A)** Time-lapse FITC fluorescence micrographs of MDCK cells under FUS treatment (*U* = 40 V, continuous waves at 1 MHz). Images shown in the three rows represent the formation of subcellular cavitation bubbles in a single cell (zoom-in 1), three different cells (zoon-in 2), and multiple cells (zoom-out), respectively. Live dye is used on cells, and the disappearance of the fluorescent signal reflects cell ablation. Arrowheads indicate the subcellular cavitation bubbles. Similar results were observed in *n* = 10 independent experiments. Scale bar: 50 μm. **(B)** Plots showing the change of live-dye fluorescence intensity within the cell cytoplasm (excluding the bubble area) over time for the cell indicated by the arrowhead in the first row in (A). **(C)** Plot of time duration from the moment of bubble formation to the moment when fluorescence intensity of the cell dropped to the background level (box: 25% - 75%, bar-in-box: median, and whiskers: min and max). *n* = 3 independent experiments; *n*_cell_ = 350. **(D)** Reconstructed 3D *z*-stack FITC fluorescence images showing subcellular bubbles generated within a GFP-expressing cell during FUS treatment. The *x*-*y*, *x*-*z*, and *y*-*z* section images were all shown. The fluorescence-depleted domain appearing in the GFP-expressing cell represents the subcellular cavitation bubble, which is indicated by the white arrowhead. Scale bar: 10 μm. **(E)** Analysis of relative fluorescence intensity at the location of the cavitation bubble. The fluorescence intensity at the center of the bubble is denoted as *h*_1_ when it first appeared, *h*_0_ before the bubble formed at the same location, and *h*_2_ after the bubble caused cell ablation, escaped, and expanded to the space above neighboring cells. Data were plotted as mean ± s.d. *n* = 3 independent experiments; *n*_cell_ = 59. **(F)** Bright field and FITC fluorescence micrographs showing subcellular cavitation bubbles generated underneath the cell and eventually causing cell membrane rupture (indicated by arrowheads) during FUS treatment. Similar results were seen in *n* = 3 independent experiments. Membrane-GFP MDCK cells were used. Scale bar: 10 μm. **(G)** Time-lapse FITC fluorescence micrographs showing bubbles appearing and disappearing underneath the cell during FUS treatment. Live dye is used to the cells. Scale bar: 10 μm. **(H)** Schematics showing different types of interactions between FUS-induced subcellular cavitation bubbles and the cell, including the unique type of cellular mechanolysis by subcellular cavitation bubbles stably growing under confined contact with the cell.

To investigate the mechanism by which bubbles induce cell ablation, we first examined the relative positioning of bubbles and cells. We performed optical sections and 3D reconstruction of GFP-expressing MDCK cells during FUS treatment. Specifically, 10 optical sections (*x*-*y* planes) were acquired along the *z*-axis, with section spacing of 11.29 μm, and the total scanning height 101.62 μm. The lowest optical section was positioned below the bottom surface of the cell (where there is no fluorescence), and the top optical section was positioned above the top surface of the cell (where there is no fluorescence) to ensure that the 3D reconstruction included the signals across entire thickness of the cell. 3D reconstructed images (**Figure 5D**) showed that the green fluorescence was uniformly distributed throughout the cell before bubble formation. As the bubble formed, a fluorescently-depleted circular domain appeared inside the cell (as shown in the *x*-*y* cross-section). Images from *x*-*z* and *y*-*z* sections further revealed that the formation of such subcellular cavitation bubble created a fluorescently-depleted volume between the cell and the substrate. These data suggest that the subcellular cavitation bubble, when growing under confined contact with the cell, might push part of the cell away from the substrate upon its formation. This suggests that under ultrasound treatment, subcellular cavitation bubbles may originate between the cell and the substrate, exerting a repulsive force on the cell.

To further confirm that abovementioned cavitation bubbles, whose appearance resulted in a fluorescently-depleted region inside the cell, were indeed formed between the cell and the substrate undeath it, we compared them with another type of bubbles that are known to be positioned above, instead of underneath, the cell, and thus do not generate a repulsive effect. Specifically, quantitative analysis showed that bubbles that eventually caused cell ablation significantly reduced the fluorescence signal at where they appear, even before the cells died (**Figure 5E**). By continuously tracking these bubbles after they ablated the cell at its original location, we observed that they remain attach to the substrate, grow, and reach the culture space above surrounding live cells. In this case, these bubbles were simply positioned above, instead of intruding into, the cells below, and these bubbles did not significantly affect the fluorescence signal from cells underneath (**Figure 5E, Figure S7A**). The contrast between above two scenarios suggests that bubbles causing cell ablation and those floating on the cell surface exhibit significant differences in optical effects, with the latter lacking a repulsive effect on the cell. This further supports our hypothesis that cavitation bubbles causing cell ablation originate between the cell and the substrate, creating repulsive forces that might eventually rupture the cell.

To further substantiate above hypothesis, we investigated the relationship between subcellular cavitation bubble formation and cell membrane deformation, by performing *z*-stack imaging on MDCK cells that only express membrane-GFP during FUS treatment (**Figure 5F**). To confirm that bubbles only appeared between the cell and the substrate, we scanned only the bottom half of the cell close to the substrate. The scanning height was 16.60 μm, consisting of 5 sections with a spacing of 4.15 μm. Under this setting, the uppermost optical section was at the middle of the cell, and the lowest was below the cell membrane, thus could capture the formation and growth of the bubble along with cell membrane deformation once the bubble formed between the cell and the substrate. Upon FUS treatment, our result showed that a subcellular cavitation bubble appeared at 7.6 s and the membrane-GFP signal enveloping the bubble was also clearly observed (Figure 5F), suggesting that the bubble indeed formed between the cell and the substrate and caused close contact with cell membrane. As the bubble gradually expanded, the membrane-GFP signal enveloping it also expanded, reflecting the direct mechanical repulsive effect of the bubble on the cell membrane. Importantly, at 38 s, the membrane-GFP signal enveloping the bubble no longer exhibited a continuous curved shape but instead showed structures resembling bifurcated cracks, followed by cell membrane rupture. This provided direct experimental evidence that the formation and growth of subcellular cavitation bubbles caused the expansion and ultimate rupture of the cell membrane (Figure 5F). Additionally, we observed that some bubbles disappeared on their own after formation, and over time, the lost fluorescence signal in the bubble-depleted domain gradually recovered. This further supports the repulsive effect caused by bubble to the cell (**Figure 5G**).

Altogether, while we do not rule out the possibility that thermal effects by FUS may contribute to the ablation of some cells within the focal zone, the above results clearly show the formation of subcellular bubbles between the cell and the substrate. Such subcellular cavitation bubbles growing under confined contact with cells impose significant mechanical loads on the cell membrane, which leads to expansion and rupture of the plasma membrane. This process deploys very rapidly (in a timeframe of seconds) and is distinct from classical mechanisms of ultrasound-induced biological effect and cell ablation, thus represents a previously unappreciated mechanics-induced mechanism, which we termed ultrasound-inflicted cellular “mechanolysis”.

Furthermore, whether bubbles exert mechanical effects on the cell nucleus is another important aspect to be examined. Therefore, we conducted FUS stimulation experiments on H2B-RFP MDCK cells, together with Live dye to monitor cell viability. The results showed that bubbles generally did not form directly beneath the cell nucleus (**Figure S7B**), and the morphology of the cell nuclei in the PAPL did not show significant distortions after cell ablation, suggesting less significant effect of subcellular bubbles on cell nucleus than that on the cell membrane. In some cells, it was indeed observed that when a bubble formed near the cell nucleus, it could cause mechanical compression and deformation of the cell nucleus, but not to a degree significant enough to induce notable translocation or rupture of the cell nucleus (**Figure S7C**). This suggests that the mechanical effect of bubbles on the cell membrane is a critical factor in FUS-inflicted cellular mechanolysis

Above findings suggest that the interactions between FUS-induced cavitation bubbles and the cell could be largely grouped into three different types (**Figure 5H**). When bubbles form only transiently before disappear between the cell and the substrate, the cell could gradually refill the depleted space created by the bubble. For bubbles that form stably and consistently grow between the substate and the cell, they cause significant deformation of the cell membrane, ultimately leading to mechanolysis of the cell. Finally, as these bubbles further expand, they may eventually penetrate through the PAPL and escape from the substrate.

To investigate whether this ultrasound-induced bubble generation can be achieved in thicker tissues, we performed FUS treatment (continuous wave) to 3D extracellular matrix made of Matrigel domes attached to the ECCD. As shown in **Figure S8A**, bubbles are continuously generated and stably expanded in the 3D matrix during the FUS treatment. *z*-stack scanning images shown in **Figure S8B** indicated that bubbles are distributed throughout the 3D matrix, indicating that this FUS-induced bubble phenomenon is not limited to monolayer cultured cells in vitro, but also occurs in more physiological-like 3D matrix.

### 2.6. Collective Cell Migration and Wound Healing after Ultrasound-Induced Cell Ablation

The wound healing response after ultrasound-induced cell ablation holds significant practical implications, yet there is a lack of exploration in previous studies. Using only mechanical scratch or place-holders to create empty space that cells could migrate into does not recapitulate key features of ultrasound-inflicted wounding, such as the residue of cell/tissue debris left at the original place after ultrasound ablation, thereby limiting their applications in understanding subsequent wound healing response. In this study, the FUS-induced ablation successfully created a unique type of wound, termed the PAPL, within a continuous epithelial monolayer, providing a more biomimetic model of ultrasound-inflicted wounding. Herein we apply this model to study the collective cell migration and wound healing after ultrasound-induced ablation.

To this end, we continued to culture cells in ECCD after ultrasound ablation and tracked the migration of the cells close to the wounded PAPL area (**Figure 6**). Under experimental conditions that allow the PAPL to remain entirely or partially attached to the substrate after ablation, we explored how it might regulate the collective migration of the neighboring live cells and the closure of the wound. Using H2B-mCherry MDCK cell monolayers, we found that live cells actively and directionally extended and migrated towards the area where the PAPL remained after ablation until the ultrasound-inflicted wound was completely closed. During this process, there remained a connection between the PAPL and live cells, and live cells also reconstructed the PAPL during the closure of the wound. Additionally, under conditions where ultrasound was strong enough to cause the detachment of the PAPL, we observed that live cells around the wound area could still migrate into the clear area left at the original place of the PAPL. This indicates that cells are intrinsically able to migrate towards the wound area regardless of the presence of the PAPL. Importantly, during the process of collective cell migration and wound healing, we observed a phenomenon of “spontaneous fracturing” between some “leading cells” at the migration front and the cells behind them, forming new tissue gaps. This implies a process of secondary damage that is generated spontaneously during the repair of the primary wound inflicted by ultrasound ablation) (**Figure S9**). This may be due to the different directions and speeds of movement of cells within the cell monolayer nearby the wound. Such secondary damages were often repaired again after the closure of the primary ultrasound-induced wound.

**Figure 6.**
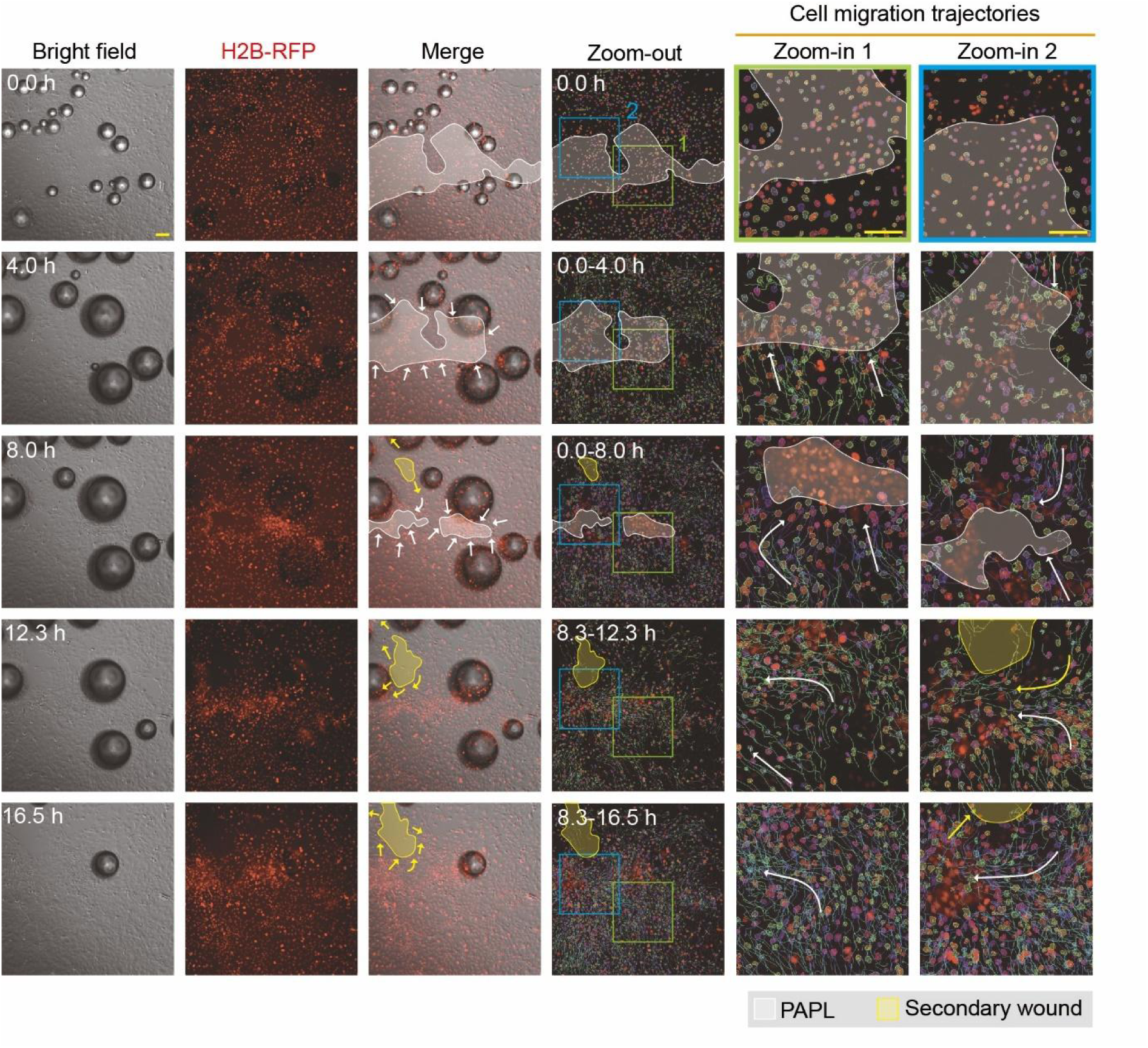
Observation of collective cell migration and wound healing after FUS-induced ablation. The first three columns display time-lapse bright-field and RFP fluorescence micrographs showing the collective migration of living H2B-RFP MDCK cells towards the PAPL area after FUS-induced ablation. In the third column, the white area represents the PAPL (with remaining RFP-labeled cell nuclei that showed no movement), with white arrows indicating the direction of collective cell migration. The yellow area represents secondary tissue gaps that appear during the primary wound healing process, largely due to the different directions and speeds of cell migration within the living cell monolayer, with yellow arrows indicating the cell migration direction. The fourth to sixth columns show the trajectories of representative cells during the collective wound healing. Colored lines represent the migration trajectory of each individual cells. The fifth to sixth columns show zoom-in views of the boxed regions. Similar results were seen in *n* = 3 independent experiments. Scale bar: 100 μm.

To further investigate the influence of the PAPL on the dynamics of collective cell migration and wound healing response after ultrasound ablation, we studied the conditions where part of the PAPL remained attached to the substrate, and tracked and analyzed the migration of individual cells nearby. As shown in **Figure 7A**, the live cell monolayer could migrate into both the PAPL and the free area (where the PAPL had detached but still available for live cells to migrate into), providing a basis for analyzing and comparing the differential guiding effect imposed by the PAPL and the free area on cell migration. As shown in **Figure 7B&C**, by tracking cell migration trajectories, we found that although the initial migration direction was relatively uniform, it was significantly influenced by the PAPL and changed direction gradually, preferentially migrating into the area where the PAPL remained. By calculating the inclined angle *θ*_n_ (counterclockwise) between the migration direction of each cell and the horizontal direction of the viewport, and comparing it with the orientational angle of the interface between the PAPL and the free area (*θ*_1_ and *θ*_2_ in Figure 7C), we found that the collective wound healing proceeded with convergent movement towards the PAPL, as reflected by decreased difference between *θ*_n_ and *θ*_1_ or *θ*_2_ over time (**Figure 7D**). In the meantime, analysis of cell movement revealed an average speed of 18 – 43.2 μm h^−1^ during the collective cell migration and wound healing process (**Figure S10**). Above results indicates that the cell migration direction gradually turned towards the PAPL, and away from the clear area, in a manner approximately aligned with the interface between the PAPL and the free area. These results suggest that the PAPL, although no longer possessing live-cell activity, still exhibits biological activity and could interact with neighboring cells to regulate their movement. This is different from the traditional free-boundary wound healing process and indicates a previously underexplored mechanism that might play an important role in dictating collective cell migration and wound healing after ultrasound-induced cell ablation. Such mechanism poses an interesting question worthy of future studies.

**Figure 7.**
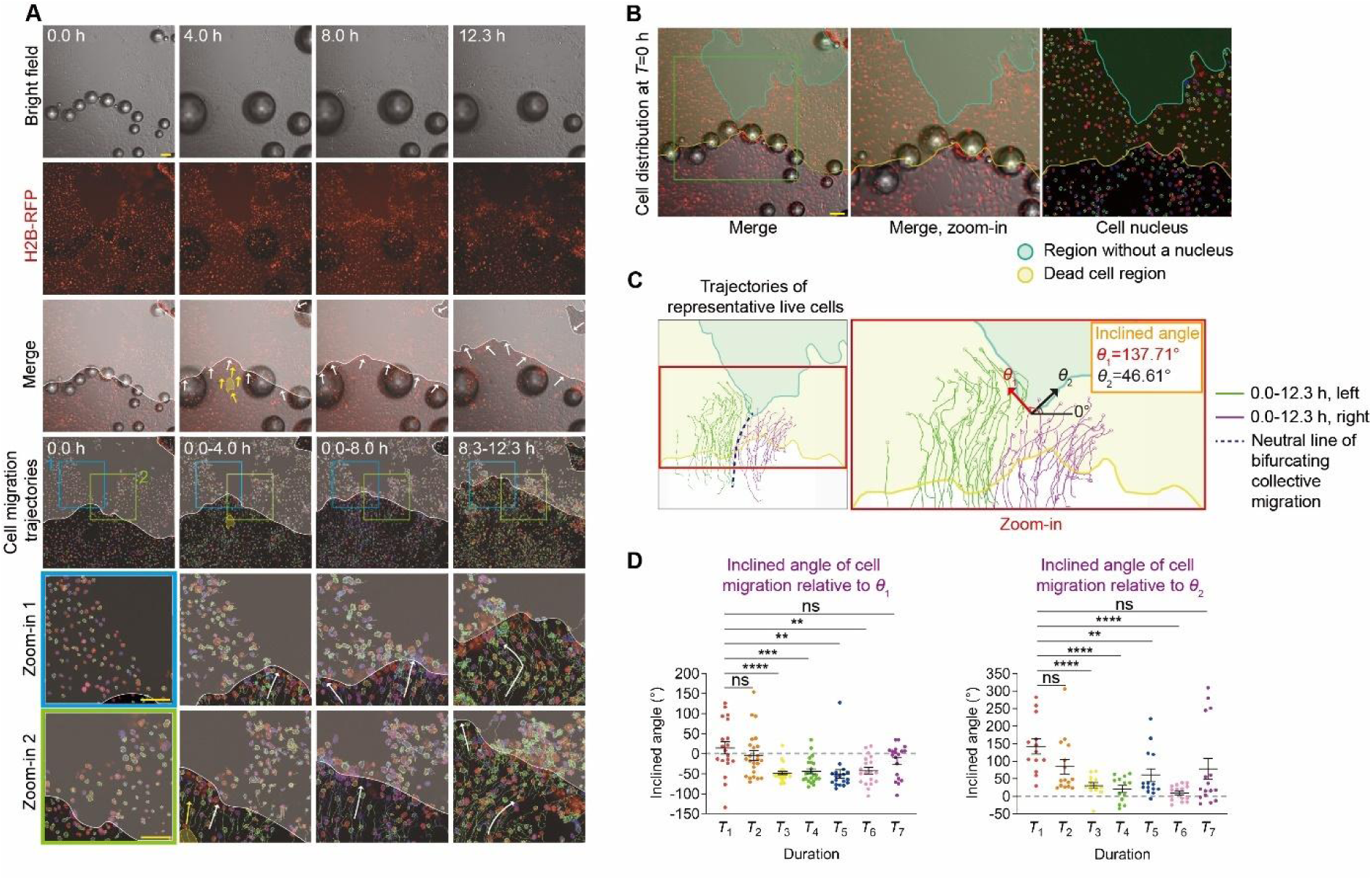
Effect of the PAPL on the collective migration and wound healing after FUS-induced ablation. **(A)** Time-lapse bright-field and RFP fluorescence micrographs showing the cell migration after FUS-induced cell ablation, featuring collective migration of living cells toward the PAPL (with remaining RFP-labeled cell nuclei that showed no movement) or the free area (where the PAPL has detached and left a nuclei-free region). The whitened area represents the PAPL or the free area, with white arrows indicating the direction of collective cell migration. The yellow area represents secondary gaps formed in the collectively migrating cell sheet nearby the wound. Yellow arrows indicate the cell migration direction near the edge of the free area. The trajectories of individual cell nuclei during the wound healing process are shown as colored lines. Blue and green boxes mark the region shown in zoom-in images. Similar results were seen in *n* = 3 independent experiments. Scale bar: 100 μm. **(B)** Confocal micrographs showing the distribution of individual cell nuclei at the beginning of collective migration. Scale bar: 100 μm. **(C)** Trajectories of individual living cells during the collective wound healing after FUS-induced ablation, which features a bifurcating migration path that is guided by the presence of PAPL flanking the clear area. The dashed line marks the neural line between the two cell groups that migrate towards the left (trajectories marked by green lines) and right (trajectories marked by purple lines) flanks of the PAPL, respectively. Similar results were seen in *n* = 3 independent experiments. *θ*_1_ and *θ*_2_ represent the orientation angle of the interface between the PAPL and the free area, which are defined and calculated as indicated in the graph. **(D)** Plots showing the inclined angle between cell migration direction and the orientation angles of the interfaces between the PAPL and the free area shown in (C). The inclined angles were calculated for the cell groups migrating towards the left (trajectories marked by green lines in (C), shown in the left plot here) and the right (trajectories marked by purple lines in (C), shown in the right plot here) flanks of the PAPL, respectively, during time duration as indicated (*T*_1_: 0.00-0.67 h; *T*_2_: 2.33-3.00 h; *T*_3_: 4.67-5.33 h; *T*_4_: 7.00-7.67 h; *T*_5_: 8.50-9.17 h; *T*_6_: 10.17-10.83 h; *T*_7_: 11.83-12.50 h). Data were plotted as mean ± s.e.m. *n*_cell_ = 34 for each indicated condition and time duration. Two-tailed Student’s *t*-test was applied to analyze the statistical significance. * *P* < 0.05, ** *P* < 0.01, *** *P* < 0.001, **** *P* < 0.0001.

## 3. Discussion

In this study, we designed and built an enclosed cell culture device capable of simultaneously achieving non-invasive ultrasound stimulation and real-time, on-the-fly live-cell imaging. Additionally, we developed an integrated platform to explore the biological effects of FUS for both short and long term in vitro. In comparison to previous experimental methods, this platform offers several advantages. Firstly, compared to conventional upright FUS-imaging systems, the design of ECCD with 45-deg ultrasound incidence angle allows a clear and uninterrupted acoustic path, thus enabling controllable FUS stimulation and ablation of cells within a pre-defined target region, while cells outside the focal area remain live. This provides a technical basis for investigating cellular responses to different levels of FUS stimulation during and after ablative treatments. Secondly, this platform is compatible with long-term cell culture and real-time microscopic imaging, wherein multiple serial excitation and continuous imaging experiments could be performed simultaneously. Thirdly, conventional upright imaging systems expose the culture system to open environment and significantly increase the risk of contamination of the culture. Even for short-term ultrasound stimulation, using upright imaging system could lead to contamination observable within 24-48 h. By avoiding cell contamination risks associated with previous methods, the ECCD platform allows the study of both the biomechanical response of cells to short-term FUS stimulation and the continuous observation of the reactive post-stimulation response of cells over an extended period. Compared to conventional systems wherein FUS treatment and real-time imaging are carried out using decoupled, open-environment devices, which often result in discontinuous and unstable experimental conditions, the ECCD platform developed in this study provides an integrative technical platform for on-demand spatiotemporal modulation of FUS treatment and continuous live-imaging in a single, enclosed, stable environment, thereby facilitating the study of biological effects and responses after ultrasound therapy.

In previous studies, cavitation effects (including stable cavitation and inertial cavitation), is the most well-accepted mechanism of FUS.^[40]^ Stable cavitation is generally triggered by low intensity ultrasound, characterized by alternating positive and negative pressures that cause stable volume oscillations around the equilibrium position of the bubble. This could exert cyclic stress and shear forces on the cell, resulting in repairable, reversible perforation.^[30,41,42]^ When the ultrasonic intensity exceeds a certain threshold, inertial cavitation causes violent volume oscillation and eventually lead to bubble rupture and collapse, producing high temperature and high pressure along with strong shock wave, microjet, or reactive oxygen (ROS), leading to reversible or irreversible perforation of cell membrane.^[43–45]^ Nevertheless, direct experimental observation of cavitation bubbles’ effect on cell ablation are still limited. In this study, our findings showed that significant cavitation bubbles could occur at the initiation site at the periphery of the focal area in the ECCD, which grow and progress before eventually leading to rapid cell death and even the breakdown of the PAPL, thereby providing further processive understanding of the effects of FUS.

Despite various types of FUS-related effects reported in previous studies, they were acquired separately from different systems that vary in the spatial and temporal scales, as well as in the absence of an integrative view of these effects at the focal region of FUS. By far, a single experimental system to study and reconcile such diverse FUS-induced phenomena remains elusive. In this work, our ECCD platform achieves real-time recording of the entire focal region under FUS treatment, which serves as an integrative system for studying the location-specific, mechanistically distinct effects throughout the focal region. Indeed, we have observed a variety of effects in the focal region, including acoustic streaming, cavitation, monolayer cracking and thermal ablation, which extends beyond classical understanding of FUS effect. This platform not only establishes a convenient system for the study of these effects, but also provides useful information for rational optimization of FUS device in the future. Particularly, in addition to classical phenomena, our results provide direct experimental evidence for the cellular mechanolysis induced by subcellular cavitation bubbles growing under confined contact with cells, which is a previously unreported biomechanical mechanism of ultrasound-induced cell ablation that occurs within a specific domain. Under moderate ultrasound intensity, these subcellular cavitation bubbles could cause mechanolysis of cells without collapse, thereby suggesting a mechanism different from that of previous studies on stable or inertial cavitation. Such cellular mechanolysis could also extend to other scenarios of applications wherein bubbles can be elicited through assistive mechanisms under LIFU. At present, in the field of LIFU combined with microbubbles, injection microbubbles are mostly used to provide cavitation nuclei, which has the advantage that the addition of injection microbubbles can significantly reduce the cavitation threshold (about 100 times), and reduce the additional damage caused by high-intensity stimulation to tissues (LIFU intensity is 0.1-1 kW cm^−2^).^[46–48]^ Together, the mechanism of cellular mechanolysis mediated by subcellular cavitation bubble formation and growth identified in this study provides a previously unrecognized principle that could operate under acoustic intensity lower than that of conventional HIFU, but still with significant cell-ablative performance, thus establishing theoretical guidance for the development of a new mode for effective, and safer, application of FUS for ablative treatments in the future.

The ECCD platform developed in this study also provides a foundation for exploring long-term biological responses following ultrasound stimulation, such as the collective wound healing after FUS-induced ablation. Currently, in vitro wound healing models are mostly based on mechanical scratches or pre-occupied void area to create free spaces for cells to migrate into.^[49,50]^ In addition to scratching, various methods such as stamping, thermal stimulation, electric shocks, and laser illumination have been employed in monolayer cell cultures to create “wounds”.^[51–54]^ However, these models remain difficult to recapitulate the tissue damage inflicted by ultrasound ablation, thereby hindering the understanding of collective cell migration during post-ablation wound healing. In this study, we achieved focal ablation in a cell monolayer and observed a residual layer of cell remnants (the PAPL) left at the site of the ablation, providing a unique model studying the collective cell migration and wound healing response after ultrasound treatment. Compared to traditional models of free-edge cell migration, our model provides a more biomimetic representation of the ultrasound-inflicted wound. The interactions between the PAPL and neighboring live cells further demonstrate its biological activity and effects that guide the direction of collective cell migration, a phenomenon not reported in previous studies. Such coupling between the PAPL and collective cell migration in wound healing are attributed to intercellular connections between live cells and the PAPL. Additionally, the release of lysophosphatidic acid during cell lysis and its activation of various downstream signals might also contribute to this phenomenon.^[55,56]^ The specific mechanisms behind this phenomenon await further exploration. In the future, this wound healing model could be used to explore the therapeutic effects of FUS-based tumor ablation and the subsequent regulatory mechanism of post-ablation wound healing process in adjacent normal tissues.

In summary, this study developed an integrated platform for real-time, on-the-fly imaging of FUS-induced cell ablation and response processes at cellular and subcellular levels, providing a unique technological basis for in-depth exploration of the mechanisms underlying ultrasound-induced biological effects. Based on this, we discovered that ultrasound could induce cellular mechanolysis through previously unappreciated subcellular cavitation bubbles that initiate and grow under confined contact with cells, adding another layer of our understanding of the biomechanical principles of ultrasound therapy. The post-ablation wound healing model further revealed that a PAPL layer could serve as a guiding cue for collective wound healing after ultrasound-inflicted ablation, offering new perspectives and methods for investigating ultrasound-based damage and reactive responses. Together, the findings of this study could help broaden our understanding of the biological effects of ultrasound, and inspire advancements in the application of FUS-based therapy.

## 4. Experimental Section

### Fabrication of enclosed cell culture device (ECCD) and transducer coupler

Firstly, high-precision 3D printing technology was utilized to print the main body of the ECCD and the coupler body using MED610 photopolymer resin (Stratasys, J750) and ABS photopolymer resin, respectively. Next, an acoustically transparent material was prepared. Mix the base prepolymer of PDMS (Dow, 184 Silicone Elastomer Kit) and the curing agent with a mass ratio of 4:1, before degassing in a vacuum chamber for 20 min. The degassed mixture is then injected with a syringe into a stainless steel mold to make 1 mm thick PDMS sheet. After further degassing for another 20 min, the mold was baked in an oven at 60 °C for 12 h. The cured PDMS sheet was removed from the mold and shaped into circular or rectangular pieces corresponding to the size of the acoustic window of the coupler or the ECCD.

The assembly of the ECCD was performed by sealing the acoustic windows with above-made PDMS sheets through the assistance of air plasma and medical silicone adhesive (Valigoo, E50, China). Similarly, a coverslip was sealed to the bottom of the ECCD for cell culture. The assembled ECCD was then dried for 12 h to allow the formation of water-light sealing that is robust even under FUS treatment. Of note, during above assembly, plasma treatment of the acoustic window materials enhanced its hydrophilicity and made it easier to remove air bubbles adhering to the inner surface of the acoustic window during cell culture. The air plasma treatment also helped reduce the adhesion of cavitation bubbles to the acoustic windows during FUS stimulation, thus preventing the bubbles from obstructing the acoustic path of ultrasound waves. The assembly of the transducer and couples was also performed similar to above. Before cell culture, ECCD was submerged in 75% ethanol for 1 h for disinfection.

### Simulation of the acoustic field in the ECCD

To verify whether the acoustic field in the ECCD meets our expectations, simulations were performed using the multiphysics simulation software COMSOL (COMSOL Multiphysics 6.1). Specifically, the “Pressure Acoustics, Frequency Domain” interface was used to calculate the acoustic pressure field, sound pressure level field, and sound intensity field when the acoustic wave stably propagates in the calculation domain under static conditions. This interface is suitable for all frequency domain simulations where the pressure field exhibits harmonic variations. To save computational resources while fully characterizing the structure of the acoustic field within the ECCD, a two-dimensional calculation domain was used. The acoustic parameters corresponding to each material in the simulation domain were assigned to the respective regions (Figure S1A&B). Regarding boundary conditions, hard acoustic boundaries (where sound waves are completely reflected) are used, except for the surface that represents the piezoelectric activator of the transducer, as well as the sound-absorbing material at the exit acoustic window. In particular, “normal displacement” condition was applied to the concave surface of the piezoelectric activator of the transducer, so as to simulate its harmonic motion along the normal direction during the generation of ultrasound at given frequency and amplitude. In addition, the “perfectly matched layer” condition was applied to the sound-absorbing material in our simulation, so that the incident acoustic waves are completely absorbed without reflection. Free triangle mesh was applied to all parts of the model except for the perfectly matched layer of the sound-absorbing material, which was set as mapped mesh. All simulations were performed under the ultrasound frequency of 1 MHz, which is consistent with the central frequency of the FUS transducer used in the experiments of this work.

### Two-level displacement control system

The entire displacement control system for the ultrasound stimulation platform was constituted by a first-level and a second-level displacement system. The first-level displacement system was manufactured by 45# steel with a tri-axial displacement control resolution of 10 μm. The second-level displacement system was a unit-direction translation stage made of 45# steel with a displacement increment resolution of 5 μm. The first-level system was used to position the transducer-coupler-ECCD assembly to the field of work of the microscope, and the second-level system was used to finely adjust the relative position of the ECCD and the coupler so as to assure co-axial alignment and first contact between them.

### FUS transducer

The FUS transducer used in this device is a single-element concave spherical transducer (PA, TXH-1-75, UK) with a central frequency of 1 MHz, a focal length of 75 mm, and a diameter of 60 mm. When conducting experiments, it is necessary to connect the water circulation system to the coupler. To minimize temperature changes of the degassed water in the coupler during ultrasound transducer operation, a circulatory cooling system was connected to the inlet / outlet on the coupler. This cooling system mainly includes a peristaltic pump (Kamoer, NKCP-S10B, China) and a temperature-controlled water bath containing degassed water. Slow circulation of degassed water was applied to the coupler to stabilize the working temperature of the transducer.

### FUS stimulation experiment using ECCD

During the experiment, the ECCD was filled with culture medium and the lid and the body of the ECCD were sealed by parafilm in a biosafety cabinet to ensure leakage-free. Ultrasound coupling agent was applied to the acoustic windows of the ECCD, before an acoustically absorbent rubber was placed onto the exterior of the exit acoustic window. Pressure was applied to eliminate air between the acoustically absorbing rubber and the acoustic window. The transducer coupler was docked to the ECCD through the entrance acoustic window, and the second-level displacement system was used for fine-adjustment of ECCD position until the entrance acoustic window is in firm contact with the coupler. During this process, it is important to ensure that the ultrasound coupling agent completely fills the space between the coupler and the entrance acoustic window of the ECCD, so as to prevent any air gaps that might obstruct the path of acoustic waves.

### Characterization of the focal domain of the FUS transducer

Firstly, we used a hydrophone system (PA, UMS3, UK) to measure the sound field generated by the FUS transducer near its focal plane. After positioning the FUS transducer in a water bath maintained at a constant temperature, a burst wave with a voltage of 20 V, frequency of 1 MHz, repetition frequency of 10 kHz, and duty cycle of 30% was generated from the transducer. The resultant acoustic field centered around the focal plane of the FUS transducer was scanned by a needle hydrophone mounted on a tri-axial displacement system. The scanning step on the *z*-axis (along the transducer axis) and on the *x* and *y* axes (perpendicular to the transducer axis) is 0.5 mm. After the scan, heat maps of the sound intensity field (*I*_sppa_) in the *x*-*y* plane and *x*-*z* plane were plotted. The focal domain was defined based on such measured distribution of the acoustic field, with the point of the highest acoustic intensity considered the focal point.

### Temperature measurement using a thermistor

To determine the dynamics of temperature change in the focal region, we applied FUS to the bottom of a T25 culture flask (Corning, #430639) where a thermistor was attached. The experimental setup is shown in Figure S3B-F. First, a thin platinum wire (50 μm in diameter) was shaped into a serpentine configuration and attached to the central region (3×3 mm^2^) of the bottom of a T25 culture flask. Subsequently, affix square copper patches (3 mm in width) on both sides of the thermistor, before soldering the end of the platinum thermistor with additional connection wires on the copper patches. Then, a layer of silicone gel was applied onto the solder points to ensure insulation and waterproofing sealing. To measure the temperature-dependent change of resistance of the thermistor, it was connected to a measurement circuit that primarily consists of a power source, two series resistors, and two parallel high-precision voltage meters. The resistance of the thermistor is calculated based on the resistors and the reading of two voltmeters. The temperature-resistance relationship of the thermistor is calibrated before the experiment. To this end, the thermistor was placed in a water bath, before continuously increasing the water bath’s temperature and recording the resistance of the thermistor in the meantime. Linear fitting of the data was used to obtain the temperature-resistance relationship.

For the measurement of temperature changes in the focal domain of the FUS transducer, a specially-made system was used. Firstly, the transducer was positioned at the bottom of a water tank filled with degassed water (maintained at a constant temperature of about 28°C) and ensured that the axis of the acoustic cone is along the vertical direction. Secondly, a T25 culture flask with the thermistor attached at the bottom was secured on a tray that is maneuverable by an electric tri-axial displacement system, before being immersed in the water bath and aligned to the acoustic path of the FUS transducer. An acoustically absorbent rubber was placed on the upper surface of the culture flask to minimize ultrasound reflections. Before experiment, it is critical to ensure the absence of air bubbles at the bottom of the culture flask or between the acoustically absorbent rubber and the culture flask. During the experiment, specific wave signals were applied to control the output of the FUS transducer, while the readings from the thermistor were recorded from the two voltage meters. The ultrasound is turned off after the readings of the voltage meters become stable, while the temperature measurement is continued until the temperature in the focal domain drops back to the environmental temperature of the water bath.

### Cell culture

Madin-Daby canine kidney cells (MDCK cells; Procell, CL-0154) were used in this study. After sterilizing the ECCD as mentioned above, the coverslip at the bottom of the ECCD was coated with polylysine to facilitate cell adhesion. Prepare a 100 μg ml^−1^ poly-L-lysine solution by diluting 1 mg ml^−1^ poly-L-lysine solution (Sigma, #P8920) with 4.5 ml of HBSS balanced salt solution (Sigma, #02-018-1A) to obtain 5 ml of the coating solution. Add the solution to the ECCD and then incubate the chamber at 37°C for 3 h. Subsequently, rinse the ECCD three times with DPBS buffer (BI, #C3590-0500).

MDCK cells were cultured in a regular 60 mm culture dish and passaged when it reached approximately 90% confluence. For cell passaging, cells were dissociated with 0.25% trypsin (BI, #C3530-0500), centrifuged at 1000 rpm for 5 min, resuspended and then replated in DMEM-based culture medium, which contains 89% DMEM base (Sigma, #D6429), 10% fetal bovine serum solution (BI, #C04001-500), and 1% antibiotic-antimycotic solution (Gibco, #15240062). Cells were cultured in an incubator under 37 °C and 5% CO_2_. The culture medium was replenished every day and cells were passaged every 3 days. Cells were used for up to P80.

### Fabrication of thermochromic inks and tapes

We used thermochromic inks that appear in red, yellow, green, and purple colors under nominal color-changing temperatures at 31, 38, 45, and 65 °C, respectively. These inks become colorless at the color-changing temperatures, while they could regain color when the temperature drops below the thresholds. To generate a gradient thermochromic ink that could display shifting colors during temperature variation, we take 4, 2, 2, and 1 ml of red, yellow, green, and purple ink, respectively, and mix them thoroughly in a beaker using a glass rod in the fume hood. Such mixed ink was applied onto adhesive tapes made of inkjet sulfuric acid paper, before air-dried in the fume hood for 12 h. Calibration of the color-changing temperatures for such gradient thermochromic tape was performed using a magnetic heating device (WIGGENS, WH260-H). Specifically, the tape was gradually heated from room temperature to 70 °C while the variation of its color was recorded, so as to establish the temperature-color relationship.

### FUS treatment and live-cell imaging experiment

Firstly, MDCK cells were cultured in ECCD until 100% confluence. To prepare for experiment, remove all liquid from the ECCD and add 60 ml pre-warmed DMEM medium at 37 °C (reaching a liquid level ∼ 5 mm above the acoustic windows of the ECCD). Subsequently, add 60 μl of the Live dye (Abbkine, #KTA1001, China) to the ECCD. Seal the lid with a parafilm and incubate in a dark environment at 37 °C for 15 min. The ECCD is then mounted on the holder of the second-level displacement system before being positioned to the work range of the microscope objective. Of note, for short-term FUS treatment experiment, it was conducted at room temperature (about 20 °C), with the ultrasound system was turned on for stimulation and cell responses were recorded in real-time using a microscope. It should be noted that in the case of such short FUS stimulation (0-10 min) without the need of further culturing, serum-free DMEM medium is sufficient. Moreover, the ECCD provides sufficient volume of culture medium (60 ml) to buffer and maintain suitable pH for long-term culture (up to 2 days) in this study, without further need to add additional CO_2_ gas infusion system in the process.

### Measurement of the temperature in ECCD by thermochromic tapes

To profile the temperature field within the cell culture area of the ECCD, a thermochromic tape is applied to the coverslip at the bottom of the ECCD, ensuring that it adheres seamlessly with any air pockets between the tape and the coverslip. The ECCD is then mounted on the tray of the second-level displacement system and the photo-recording device is placed under the ECCD. While the FUS stimulation is on, the color change of the thermochromic tape is recorded in the meantime. Immediately after FUS treatment, the tape was removed from the bottom of the ECCD and the cell ablation results were observed with an inverted fluorescence microscope. A mosaic mode is adopted during image acquisition to achieve the scanning of the entire cell culture area.

### High-temperature heat bath experiment

ECCD containing MDCK cells (cultured ∼95% confluence) were used along with Live dye to monitor cell viability before and after the heat bath. ECCD was seal by parafilm and submerged in a water bath pre-set at 65 and 85 °C, respectively, for 5 min. Images of cells were acquired both before and after the heat bath to calculate the relative cell ablation area directly caused by such high-temperature heat bath.

### Preparation of 3D matrix on the bottom of ECCD

Firstly, 30 μl Matrigel matrix (Corning, 354234) was fully mixed with 30 μl PBS on ice, before the mixture was deposited to the center of the bottom of the ECCD to form a dome with a diameter of about 6 mm and a height of 4 mm. Then the ECCD was incubated at 37 °C for 20 min for matrix solidification, before additional 60 ml of PBS was added to the ECCD and stored until use.

### Cell migration trajectory analysis

The H2B-RFP MDCK cells were used to enable the tracking of cell migration. Different from the short-term FUS stimulation, the medium used for long-term cell culture was DMEM medium (60 ml per ECCD) supplemented with 10% fetal bovine serum and 1% antibiotic-antifungal medium. Due to the excessive amount of culture medium, it was sufficient to buffer and maintain stable pH during long-term cell culture and imaging in this study, without additional need to replenish the medium or infuse CO_2_. In the FUS ablation experiment for cell migration study, the voltage of the FUS transducer is 40 V, emission frequency 1 MHz (continuous wave), and treatment time 3 min. After the treatment, the FUS-treated ECCD was cultured continuously on the microscope platform for another 24 h with multi-point image acquisition applied under the bright field and RFP fluorescence channels. Time-lapse microscopic images of the migration of living cell into the post-ablation phantom layer or the cleared region (where the residual phantom layer detached) were acquired under the bright field and RFP fluorescence channels after the FUS treatment.

The 2D tracking module in the NIS-Elements AR software (Nikon, Ti2-E, Japan) was used to analyze the time-lapse images of cell migration as acquired above. This allows the tracking of individual cells within a collectively migrating cell population over a period of time. Specifically, the RFP-labeled cell nuclei were used as markers for cell migration trajectory analysis. Using the 2D tracking module, each nucleus is first identified and labeled, and then the migration trajectory of the nucleus is plotted based on the time-lapse changes of its location. Cell migration speed and step length were also analyzed using above method.

### Statistical Analysis

Samples were randomized into experimental groups. Control of covariates is not needed because pure cell lines are used and all experimental parameters were kept constant (except for those parameters specifically tested by the experiment). Investigators were blinded to group allocation during data analysis. No experimental units or data were excluded. All results were from *n* ≥ 3 independent experiments. Data were presented at mean ± s.d. or s.e.m. as indicated. Student’s *t*-test (two-tailed) was used to assess the statistical significance. GraphPad Prism 10.0.2 was used to analyze significance. Significance level was defined by *p*-value in all plots.

## Supporting Information

Supporting Information is available from the Wiley Online Library or from the author.

## Author Contributions

Z.B. and Y.S. conceived the project; Z.B. designed and performed experiments; Z.B., Z.L., and Y.S. analyzed data. Y.S. supervised the project. All authors contributed to the manuscript.

## Acknowledgements

We thank Dr. Haidong Wang for assistance in thermistor measurement. We thank Dr. Jiabin Zhang and Dr. Feng Feng for assistance in ultrasound transducer calibration. We thank Dr. Yan Yi for the gift of the ultrasound-absorbent rubber. We thank Dr. Jianbo Bai for the gift of the H2B-RFP MDCK cell line. Y.S.’s work is supported by the National Natural Science Foundation of China (U21A20203, 12102229, 11921002), the National Key Research and Development Program of China (2021YFA0719301), the Oversea High-level Scholar Introduction Program, Tsinghua University Dushi Program, and the Tsinghua University Startup Funding.

## Data Availability Statement

The data that support the findings of this study are available from the corresponding author upon reasonable request.

## Ethical Statement

We declare no ethics concerns related to this study.

## References

[1] V. Krishna, F. Sammartino, A. Rezai, JAMA Neurol. 2018, 75, 246.

[2] Z. Izadifar, Z. Izadifar, D. Chapman, P. Babyn, J. Clin. Med. 2020, 9, 460.

[3] Y. Meng, K. Hynynen, N. Lipsman, Nat. Rev. Neurol. 2020, 17, 7.

[4] F. Recker, M. Thudium, H. Strunk, T. Tonguc, S. Dohmen, G. Luechters, B. Bette, S. Welz, B. Salam, K. Wilhelm, E.K. Egger, U. Wüllner, U. Attenberger, A. Mustea, R. Conrad, M. Marinova, Sci. Rep. 2021, 11,

[5] Y. Tang, C. Chen, B. Jiang, L. Wang, F. Jiang, D. Wang, Y. Wang, H. Yang, X. Ou, Y. Du, Q. Wang, J. Zou, Int J Nanomed 2021, Volume 16, 4643.

[6] N. Lipsman, Y. Meng, A.J. Bethune, Y. Huang, B. Lam, M. Masellis, N. Herrmann, C. Heyn, I. Aubert, A. Boutet, G.S. Smith, K. Hynynen, S.E. Black, Nat. Commun. 2018, 9,

[7] S.R. Sirsi, M.A. Borden, Adv. Drug Deliver. Rev. 2014, 72, 3.

[8] K.F. Timbie, U. Afzal, A. Date, C. Zhang, J. Song, G. Wilson Miller, J.S. Suk, J. Hanes, R.J. Price, J. Control Release. 2017, 263, 120.

[9] S. Alli, C.A. Figueiredo, B. Golbourn, N. Sabha, M.Y. Wu, A. Bondoc, A. Luck, D. Coluccia, C. Maslink, C. Smith, H. Wurdak, K. Hynynen, M. O’Reilly, J.T. Rutka, J. Control Release. 2018, 281, 29.

[10] Y. Cao, Y. Chen, T. Yu, Y. Guo, F. Liu, Y. Yao, P. Li, D. Wang, Z. Wang, Y. Chen, H. Ran, Theranostics 2018, 8, 1327.

[11] M.B.C. de Matos, R. Deckers, B. van Elburg, G. Lajoinie, B.S. de Miranda, M. Versluis, R. Schiffelers, R.J. Kok, Front. Pharmacol. 2019, 10,

[12] Y. Meng, R.M. Reilly, R.C. Pezo, M. Trudeau, A. Sahgal, A. Singnurkar, J. Perry, S. Myrehaug, C.B. Pople, B. Davidson, M. Llinas, C. Hyen, Y. Huang, C. Hamani, S. Suppiah, K. Hynynen, N. Lipsman, Sci. Transl. Med. 2021, 13,

[13] M.A. Stavarache, N. Petersen, E.M. Jurgens, E.R. Milstein, Z.B. Rosenfeld, D.J. Ballon, M.G. Kaplitt, J. Neurosurg. 2019, 130, 989.

[14] N. Sheikov, N. McDannold, N. Vykhodtseva, F. Jolesz, K. Hynynen, Ultrasound Med. Biol. 2004, 30, 979.

[15] H. Chen, E.E. Konofagou, J. Cereb. Blood Flow Metab. 2014, 34, 1197.

[16] C.T. Curley, B.P. Mead, K. Negron, N. Kim, W.J. Garrison, G.W. Miller, K.M. Kingsmore, E.A. Thim, J. Song, J.M. Munson, A.L. Klibanov, J.S. Suk, J. Hanes, R.J. Price, Sci. Adv. 2020, 6,

[17] M.N. Centelles, M. Wright, P.-W. So, M. Amrahli, X.Y. Xu, J. Stebbing, A.D. Miller, W. Gedroyc, M. Thanou, J. Control Release. 2018, 280, 87.

[18] P. Cressey, M. Amrahli, P.-W. So, W. Gedroyc, M. Wright, M. Thanou, Biomaterials 2021, 271, 120758.

[19] P.C. Rinaldi, J.P. Jones, F. Reines, L.R. Price, Brain Res. 1991, 558, 36.

[20] P.-H. Tsui, S.-H. Wang, C.-C. Huang, Ultrasonics 2005, 43, 560.

[21] D.P. Darrow, P. O’Brien, T.J. Richner, T.I. Netoff, E.S. Ebbini, Brain Stimul. 2019, 12, 1439.

[22] T. Zhang, Z. Wang, H. Liang, Z. Wu, J. Li, J. Ou-Yang, X. Yang, Y.B. Peng, B. Zhu, IEEE. T. Bio-med. Eng. 2022, 69, 3155.

[23] V. Cotero, J. Graf, H. Miwa, Z. Hirschstein, K. Qanud, T.S. Huerta, N. Tai, Y. Ding, K. Jimenez-Cowell, J.N. Tomaio, W. Song, A. Devarajan, T. Tsaava, R. Madhavan, K. Wallace, E. Loghin, C. Morton, Y. Fan, T.-J. Kao, K. Akhtar, M. Damaraju, L. Barenboim, T. Maietta, J. Ashe, K.J. Tracey, T.R. Coleman, D. Di Carlo, D. Shin, S. Zanos, S.S. Chavan, R.I. Herzog, C. Puleo, Nat. Biomed. Eng. 2022, 6, 683.

[24] Y. Zhong, Y. Zhang, J. Xu, J. Zhou, J. Liu, M. Ye, L. Zhang, B. Qiao, Z.-g. Wang, H.-t. Ran, D. Guo, ACS Nano 2019, 13, 3387.

[25] D.P. Zachs, S.J. Offutt, R.S. Graham, Y. Kim, J. Mueller, J.L. Auger, N.J. Schuldt, C.R.W. Kaiser, A.P. Heiller, R. Dutta, H. Guo, J.K. Alford, B.A. Binstadt, H.H. Lim, Nat. Commun. 2019, 10,

[26] T.S. Huerta, A. Devarajan, T. Tsaava, A. Rishi, V. Cotero, C. Puleo, J. Ashe, T.R. Coleman, E.H. Chang, K.J. Tracey, S.S. Chavan, Sci. Rep. 2021, 11,

[27] J.E. Kennedy, Nat. Rev. Cancer 2005, 5, 321.

[28] H.P. Kok, E.N.K. Cressman, W. Ceelen, C.L. Brace, R. Ivkov, H. Grüll, G. ter Haar, P. Wust, J. Crezee, Int. J. Hyperther. 2020, 37, 711.

[29] K. Hynynen, R. Roemer, D. Anhalt, C. Johnson, Z.X. Xu, W. Swindell, T. Cetas, Int. J. Hyperther. 2009, 3, 21.

[30] J. Wu, W.L. Nyborg, Adv. Drug Deliver. Rev. 2008, 60, 1103.

[31] S. Hu, X. Zhang, M. Unger, I. Patties, A. Melzer, L. Landgraf, Cells 2020, 9, 2595.

[32] M. Kinoshita, K. Hynynen, Biochem. Biophys. Res. Commun. 2005, 335, 393.

[33] R.K. Schlicher, H. Radhakrishna, T.P. Tolentino, R.P. Apkarian, V. Zarnitsyn, M.R. Prausnitz, Ultrasound Med. Biol. 2006, 32, 915.

[34] Z. Qiu, J. Guo, S. Kala, J. Zhu, Q. Xian, W. Qiu, G. Li, T. Zhu, L. Meng, R. Zhang, H.C. Chan, H. Zheng, L. Sun, iScience 2019, 21, 448.

[35] Y.-S. Huang, C.-H. Fan, N. Hsu, N.-H. Chiu, C.-Y. Wu, C.-Y. Chang, B.-H. Wu, S.-R. Hong, Y.-C. Chang, A. Yan-Tang Wu, V. Guo, Y.-C. Chiang, W.-C. Hsu, L. Chen, C. Pin-Kuang Lai, C.-K. Yeh, Y.-C. Lin, Nano Lett. 2019, 20, 1089.

[36] H. Kitamori, I. Sumida, T. Tsujimoto, H. Shimamoto, S. Murakami, M. Ohki, Phys. Medica. 2019, 58, 90.

[37] Y. Zhao, J. Xu, Y. Zhang, F. Wu, W. Zhao, R. Li, Y. Yang, M. Zhang, Y. Zhang, C. Guo, Chem. Eng. J. 2023, 472, 144911.

[38] G.E. Davis, J. Opt. Soc. Am. 1955, 45, 572.

[39] P.L. Marston, Sonochemistry and Sonoluminescence 1999, 73.

[40] X. Fu, X. Hu, ACS Applied Bio Materials 2024, 7, 8040.

[41] H. Honda, Q.-L. Zhao, T. Kondo, Ultrasound Med. Biol. 2002, 28, 673.

[42] A.A. Doinikov, A. Bouakaz, J. Acoust. Soc. Am. 2010, 128, 11.

[43] M. Lokhandwalla, B. Sturtevant, Phys. Med. Biol. 2001, 46, 413.

[44] E.A. Brujan, Ultrasound Med. Biol. 2004, 30, 381.

[45] Z. Jiang, W. Xiao, Q. Fu, J. Control Release. 2023, 361, 547.

[46] J. Deprez, G. Lajoinie, Y. Engelen, S.C. De Smedt, I. Lentacker, Adv. Drug Deliver. Rev. 2021, 172, 9.

[47] A. Dauba, A. Delalande, H.A.S. Kamimura, A. Conti, B. Larrat, N. Tsapis, A. Novell, Pharmaceutics 2020, 12, 1125.

[48] M. Hoogenboom, D. Eikelenboom, M.H. den Brok, A. Heerschap, J.J. Fütterer, G.J. Adema, Ultrasound Med. Biol. 2015, 41, 1500.

[49] D. Fox, H. Tao, A.J. Berno, D.R. Cox, K.A. Frazer, PLoS One 2007, 2, e697.

[50] N.B. Wolf, S. Küchler, M.R. Radowski, T. Blaschke, K.D. Kramer, G. Weindl, B. Kleuser, R. Haag, M. Schäfer-Korting, Eur. J. Pharm. Biopharm 2009, 73, 34.

[51] A. Stamm, K. Reimers, S. Strauß, P. Vogt, T. Scheper, I. Pepelanova, BioNanoMaterials 2016, 17, 79.

[52] K. Safferling, T. Sütterlin, K. Westphal, C. Ernst, K. Breuhahn, M. James, D. Jäger, N. Halama, N. Grabe, J. Cell Biol. 2013, 203, 691.

[53] J.W.J. van Kilsdonk, E.H. van den Bogaard, P.A.M. Jansen, C. Bos, M. Bergers, J. Schalkwijk, Wound Repair Regen. 2013, 21, 890.

[54] S. Ud-Din, A. Bayat, Wound Repair Regen. 2017, 25, 164.

[55] G. Broughton, J.E. Janis, C.E. Attinger, Plast. Reconstr. Surg. 2006, 117, 12S.

[56] S. Schreml, R.-M. Szeimies, L. Prantl, M. Landthaler, P. Babilas, J. Am. Acad. Dermatol. 2010, 63, 866.

