## Supplementary Information for "Subcellular Cavitation Bubbles Induce Cellular Mechanolysis and Collective Wound Healing in Ultrasound-Inflicted Cell Ablation"

**Figure S1**

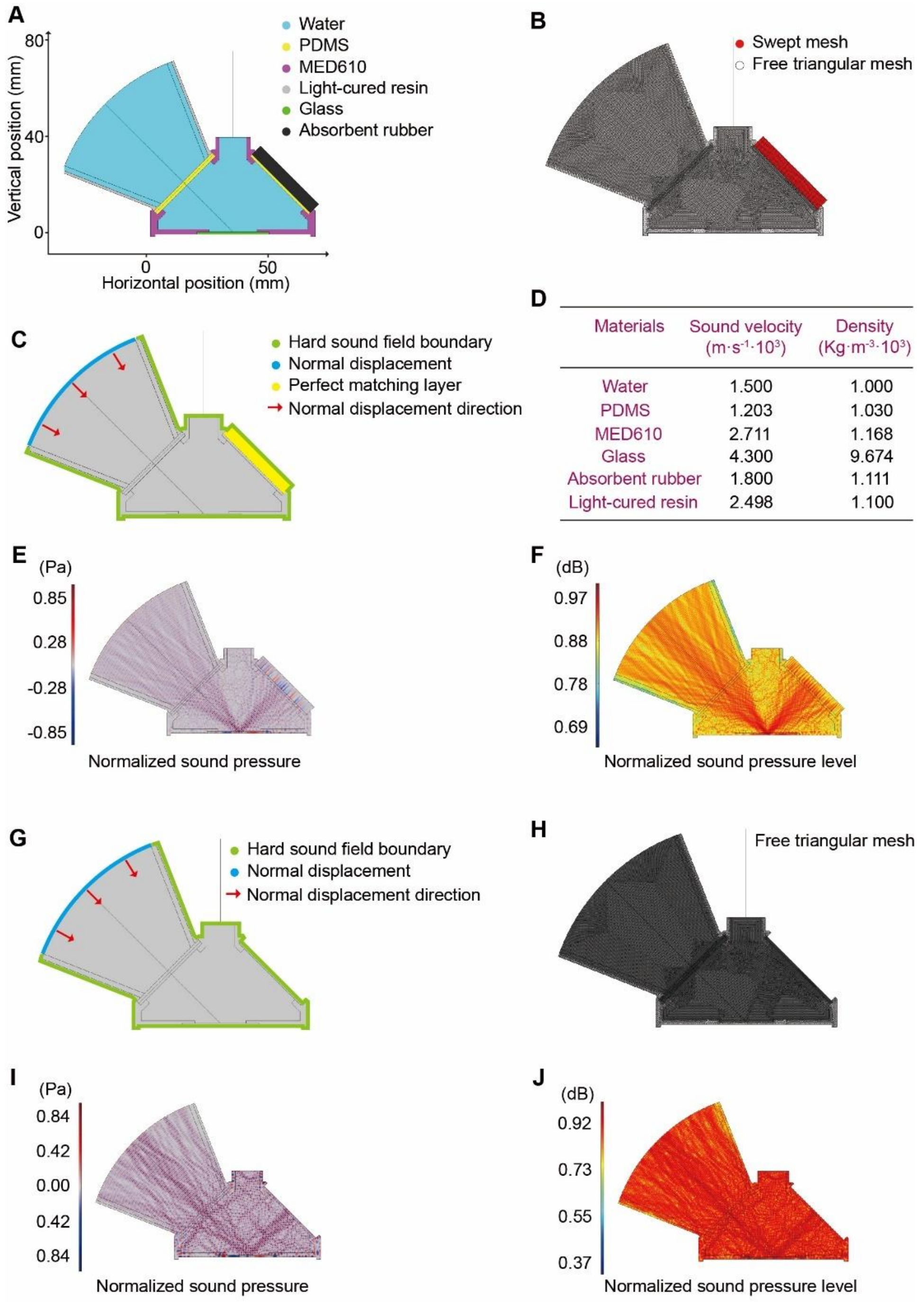


**Figure S1. Simulation of the acoustic field in the ECCD under FUS stimulation. (A-C)** Two-dimensional simulation domain of the ECCD assembled with the FUS transducer (A), with corresponding meshing (B) and boundary conditions (C) applied to the model. **(D)** Material parameters assigned to different parts of the model. **(E&F)** Simulation results of the acoustic pressure field (E) and sound pressure level field (F) within the ECCD under FUS stimulation. **(G&H)** Schematic showing the boundary conditions (G) and meshing (H) assigned to the two-dimensional simulation domain of the ECCD without sound-absorbing rubber at the exit acoustic window (ExAW). **(I&J)** Simulation results of the acoustic pressure field (I) and sound pressure level field (J) within the ECCD in the absence of sound-absorbing rubber at the ExAW.

**Figure S2**

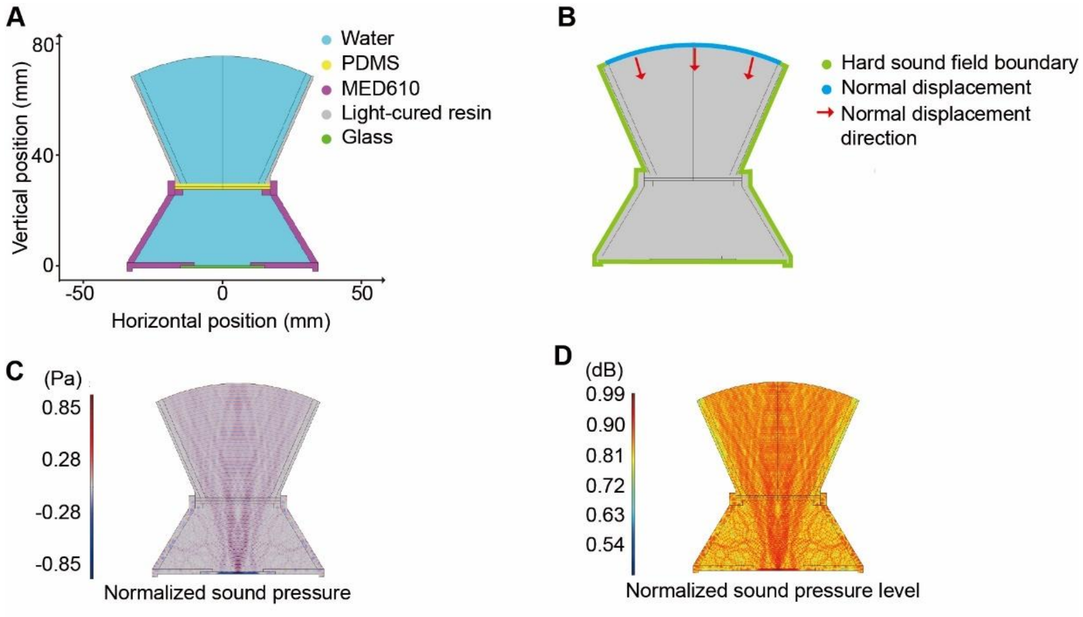


**Figure S2. Simulation of acoustic field within a conventional upright microscope-transducer system. (A&B)** 2D simulation model (A) and boundary conditions (B) of an upright microscope-transducer system combined with a cell culture device. **(C&D)** Simulation results the acoustic pressure field (C) and sound pressure level field (D) within this upright system.

### Figure S3

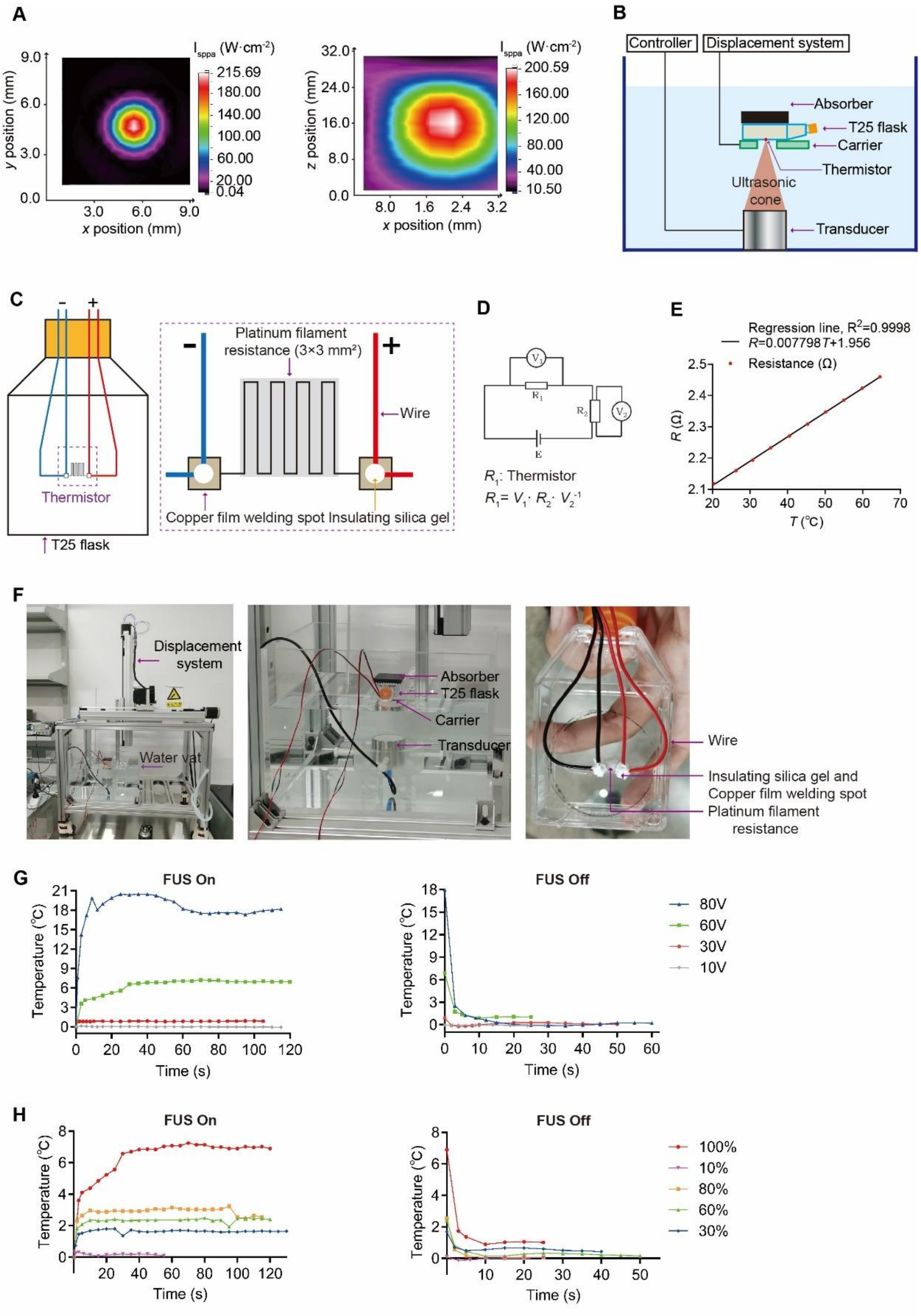


**Figure S3. Characterization of the focal region and thermal effects of the FUS transducer used in this study.** **(A)** The acoustic intensity field diagram ($I_{sppa}$) of the *x*-*y* and *x*-*z* planes of the focal region of the FUS transducer ($U$ = 20 V, FUS frequency of 1 MHz, repetition frequency of 10 kHz, 30% duty cycle). **(B)** Schematic of the experimental setup for measuring the thermal effect induced by the FUS transducer within the focal domain. The FUS is emitted upwards, and the platinum wire-based thermistor is attached to the bottom surface of the culture flask that is positioned at the focal point of the FUS transducer. **(C)** Schematic showing the thermistor at the bottom of a T25 flask and related wiring for the measurement of resistance. **(D)** Diagram of electric circuit used for the measurement of temperature-dependent changes in the resistance of the thermistor. **(E)** Calibrated temperature-resistance relationship of the platinum-wire thermistor. **(F)** Photograph of the experimental setup for measuring the thermal effect in the focal domain of the FUS transducer. **(G)** Plots showing the temperature change at the focal domain when the FUS is on and off, respectively, under indicated transducer voltage. **(H)** Plots showing the temperature change at the focal domain when the FUS is on and off, respectively, under indicated duty cycles. FUS operates with a burst wave at $U = 60$ V, a repetition frequency of 10 kHz.

**Figure S4**

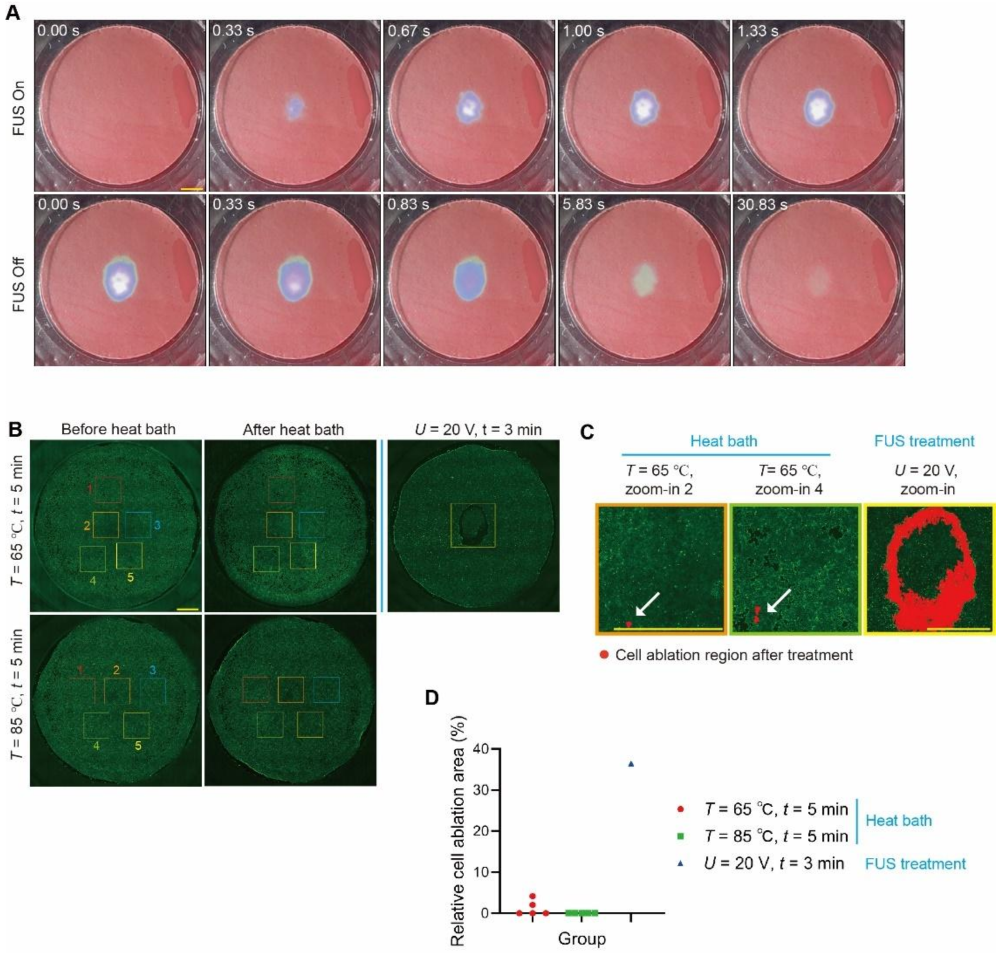


**Figure S4. Analysis of temperature field and cell field under FUS treatment. (A)** Representative images showing the time-dependent changes of the distribution of temperature within the cell culture region of the ECCD upon FUS treatment ($U$ = 50 V, continuous waves of 1 MHz, $T$ = 3 min). Scale bar: 5 mm. **(B)** The FITC fluorescence images of MDCK cells before and after the treatment with heat bath at 65 and 85 °C, respectively, for 5 min. Bottom panel shows the FITC fluorescence image of cells after FUS treatment ($U$ = 20 V, continuous waves of 1 MHz, $T$ = 3 min). Cells are labeled with Live dye. The areas boxed by colored squares were used to represent areas where the relative cell ablation area is analyzed. **(C)** Zoom-in views of boxed areas indicated in (B). The red area indicated by the white arrow is the area where the cells are ablated during the heat bath or during the FUS stimulation. Scale: 5 mm. **(D)** Plot of the relative cell ablation area under indicated conditions.

**Figure S5**

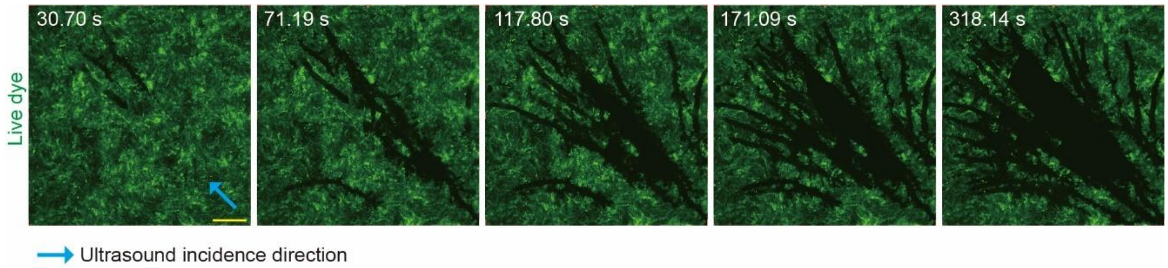


**Figure S5. FUS-induced streak-like cell ablation domains at the central area of the focal region.** Time-lapse FITC fluorescence micrograph showing the distribution of living cells in focal domain during FUS treatment ($U$ = 40 V, continuous waves of 1 MHz, $T$ = 6 min). Live cells were labeled with FITC Live dye. The white arrow indicates the direction of incidence ultrasound waves. Similar results were seen in $n$ = 3 independent experiments. Scale bar: 500 μm.

**Figure S6**

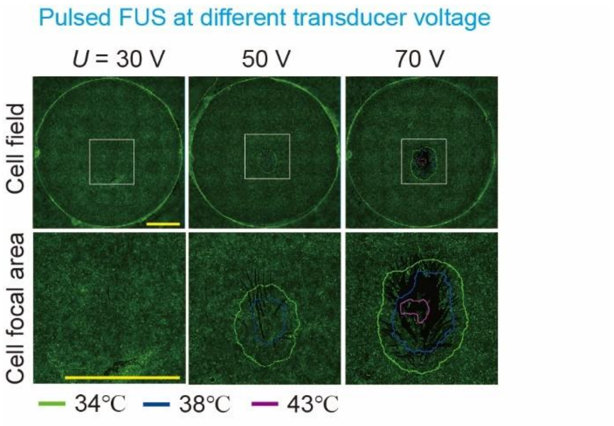


**Figure S6. Pulsed FUS did not induce cavitation bubbles and the elliptical cell ablation band.** Time-lapse FITC fluorescence micrograph showing the distribution of live cells and the corresponding temperature field in the focal domain during pulsed FUS treatment. Only streak-like ablation domains were formed without bubble generation. No notable bubble-associated elliptical ablation band was observed under pulsed FUS due to lower energy intensity. The FUS transducer operates at 40 V with a frequency of 1MHz. The pulse duty cycle is 10%, the pulse repetition rate is 1kHz, and the treatment duration is 6 min. Live cells were labeled with FITC Live dye. Scale bar: 5 mm. The cells used in the above experiments were wild-type MDCK cells.

**Figure S7**

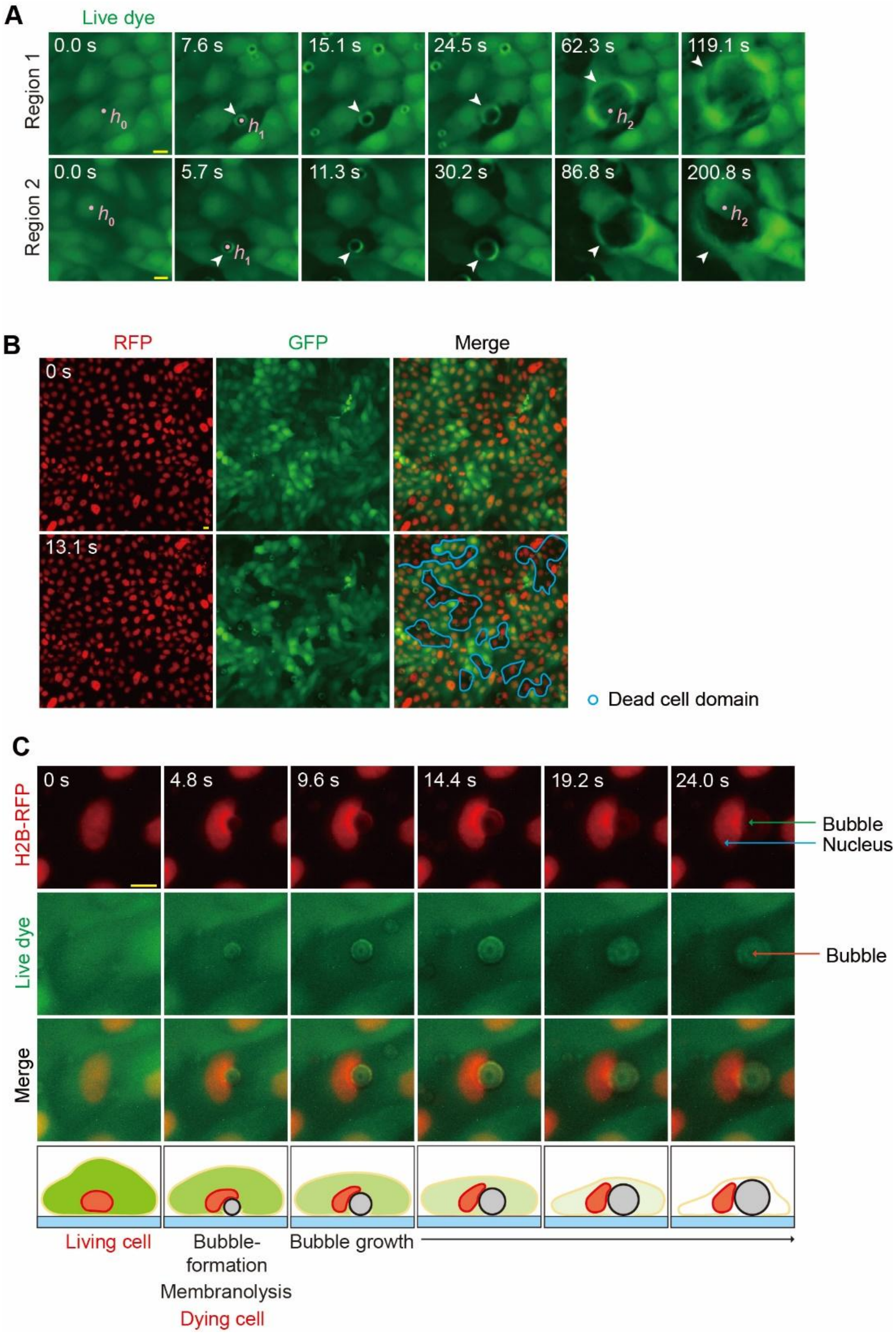


**Figure S7. Investigating the interaction between cavitation bubbles and cells. (A)** Time-lapse FITC fluorescence micrographs showing the formation of bubbles, cell ablation, and bubble growth during FUS treatment ($U$ = 40 V, continuous waves of 1 MHz). Live dye is used on cells. The dots mark the position for measuring fluorescence intensities $h_0$, $h_1$ and $h_2$ in **Figure 5E**. Scale bar: 10 μm. **(B)** Time-lapse confocal micrographs showing the change of the distribution of cell nuclei (red) and living cells (green, Live dye), respectively, during FUS treatment ($U$ = 40 V, continuous waves of 1 MHz). Cells are H2B-RFP MDCK cells, and live dye (green) is added. Scale bar: 10 µm. **(C)** Time-lapse confocal micrographs showing the presence of stably growing subcellular cavitation bubbles nearby a nucleus during FUS treatment ($U$ = 40 V, continuous waves of 1 MHz). Cells are H2B-RFP MDCK cells, and live dye is added. Scale bar: 10 µm. The bottom panel represents a schematic that summarizes the changes of the cell, cell nucleus, and cavitation bubbles over time. The green shade represents the cell, the yellow lines represent the cell membrane, the red shade represents the cell nucleus, the blue shade represents the cell culture substrate, and the gray circle represents the cavitation bubble.

**Figure S8**

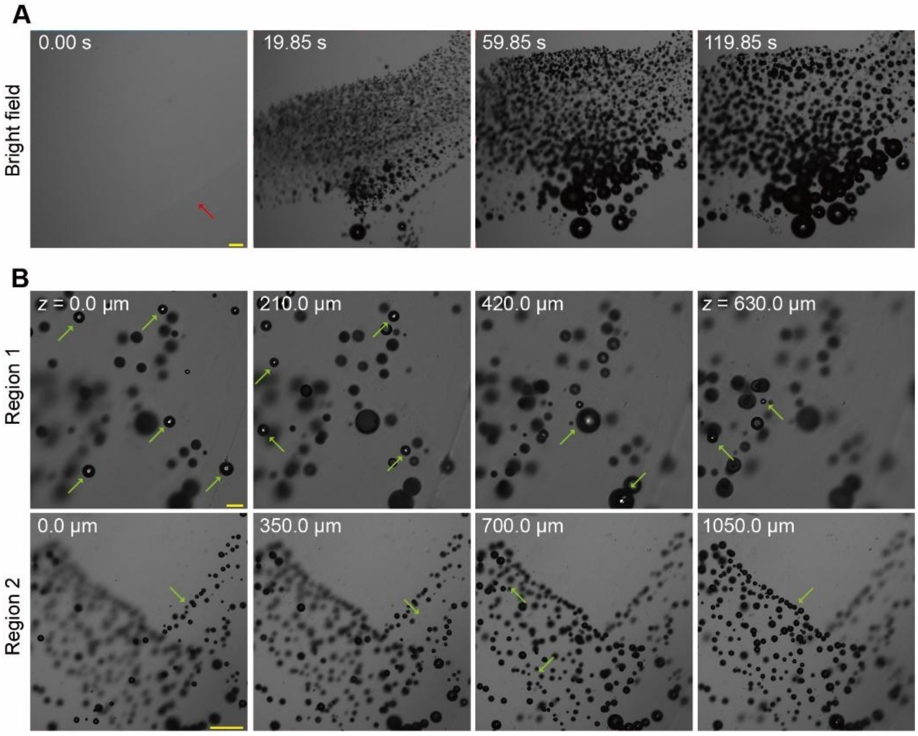


**Supplementary Figure S8. FUS-induced stable bubble formation within 3D extracellular matrix. (A)** Time-lapse confocal micrographs showing the formation and expansion of bubbles in the 3D Matrigel matrix over time during FUS treatment ($U = 40$ V, continuous waves of 1 MHz). The red arrow indicates the gel bead and its boundary. Scale :100 µm. **(B)** *Z*-stack confocal micrographs showing the formation and distribution of cavitation bubbles at different heights throughout the 3D matrix after FUS treatment ($U = 40$ V, continuous waves of 1 MHz, $T = 5$ min). Similar results were seen in $n = 3$ independent experiments. Scale bars: 100 µm (upper panels), 500 µm (lower panels).

**Figure S9**

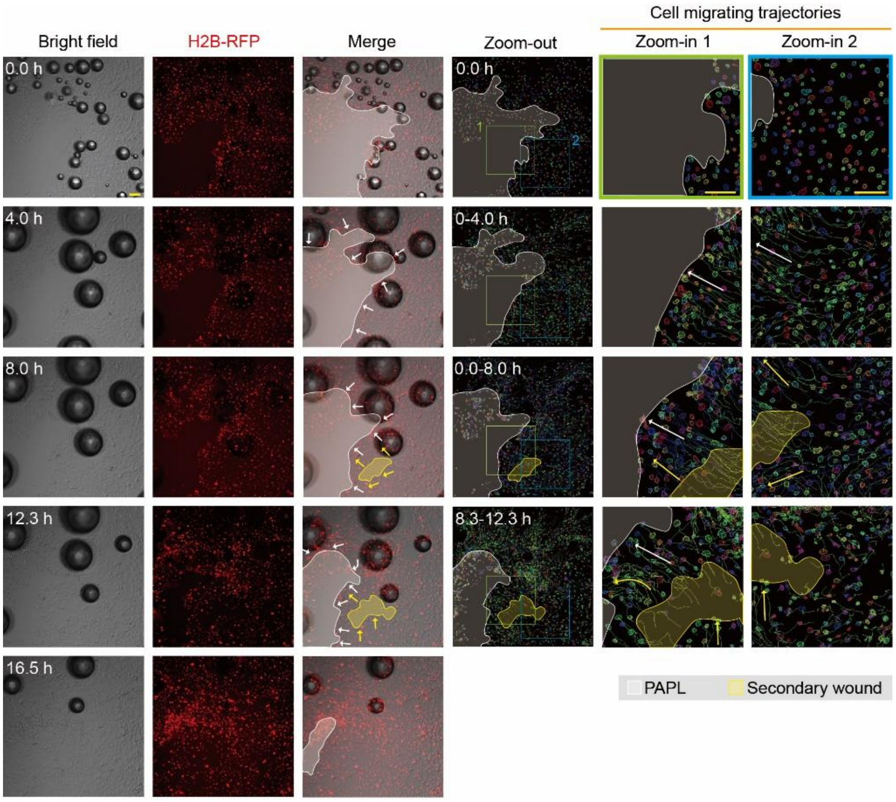


**Figure S9. Collective wound healing after FUS-induced ablation.** The first three columns display time-lapse bright-field and RFP fluorescence micrographs showing the collective migration of living cells towards the PAPL or clear area after FUS-induced ablation. In the third column, the white area represents the PAPL (with remaining RFP-labeled cell nuclei but no movement) or the clear area (where PAPL detached), with white arrows indicating the direction of collective cell migration. The yellow area represents secondary tissue gaps that appear during the primary wound healing process, with yellow arrows indicating the cell migration direction. The fourth to sixth columns show the trajectories of representative cells during the collective wound healing. Colored lines represent the migration trajectory of each individual cells. The fifth to sixth columns show zoom-in views of the boxed regions. Scale bar: 100 μm.

**Figure S10**

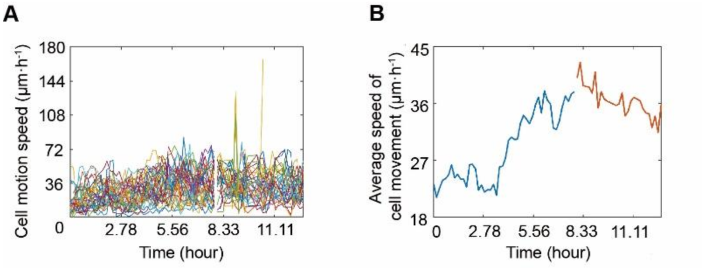


**Figure S10. Migration speed during collective wound healing after FUS-induced ablation. (A&B)** Line graphs showing the migration speed of individual cells ($n_{cell}$ = 5896) (A) and the average speed of the cell population (B) during collective migration after FUS-induced ablation. *n* = 3 independent experiments.

**Table S1.** The acoustic parameters of selected materials at environmental temperature of 25 ℃.

| Materials | Sound velocity ($10^3$ m $s^{-1}$) | Density ($10^3$ kg $m^{-3}$) | Acoustic impedance ($10^6$ kg $m^{-2}$ $s^{-1}$) |
|---|---|---|---|
| Plastic (T25 flask) | 2.766 | 1.035 | 2.863 |
| Plastic (35mm dish) | 3.020 | 1.046 | 3.159 |
| Silica gel | 1.003 | 1.049 | 1.052 |
| Acrylic plate | 4.023 | 1.189 | 4.783 |
| Plastic (culture plate) | 2.753 | 1.044 | 2.874 |
| MED610 light-cured resin | 2.711 | 1.168 | 3.166 |
| PDMS (4:1) | 1.203 | 1.030 | 1.239 |
| Pure water | 1.500 | 1.000 | 1.500 |

**Table S2.** The acoustic parameters of 1 mm thick PDMS sheet made by different mixing ratio of base pre-polymer and curing agent (environmental temperature of 25 ℃).

| Ratio of base to curing agent | Sound velocity ($10^3$ m $s^{-1}$) | Density ($10^3$ kg $m^{-3}$) | Acoustic impedance ($10^6$ kg $m^{-2}$ $s^{-1}$) |
|---|---|---|---|
| 10:1 | 2.766 | 1.035 | 2.863 |
| 5:1 | 3.020 | 1.046 | 3.159 |
| 4:1 | 1.003 | 1.049 | 1.052 |
| 3:1 | 4.023 | 1.189 | 4.783 |
| 2:1 | 2.753 | 1.044 | 2.874 |
| 1:1 | 1.052 | 1.024 | 1.077 |
